# Comparing two circular distributions: advice for effective implementation of statistical procedures in biology

**DOI:** 10.1101/2021.03.25.436932

**Authors:** Lukas Landler, Graeme D. Ruxton, E. Pascal Malkemper

## Abstract

Many biological variables, often involving timings of events or directions, are recorded on a circular rather than linear scale, and need different statistical treatment for that reason. A common question that is asked of such circular data involves comparison between two groups or treatments: Are the populations from which the two samples drawn differently distributed around the circle? For example, we might ask whether the distribution of directions from which a stalking predator approaches its prey differs between sunny and cloudy conditions; or whether the time of day of mating attempts differs between lab mice subject to one of two hormone treatments. An array of statistical approaches to these questions have been developed. We compared 18 of these (by simulation) in terms of both abilities to control type I error rate near the nominal value, and statistical power. We found that only eight tests offered good control of type I error in all our test situations. Of these eight, we are able to identify Watson’s U^2^ test and MANOVA based on trigonometric functions of the data as offering the best power in the overwhelming majority of our test circumstances. There was often little to choose between these tests in terms of power, and no situation where either of the remaining six tests offered substantially better power than either of these. Hence, we recommend the routine use of either Watson’s U^2^ test or MANOVA when comparing two samples of circular data.

## Introduction

Many variables in biology are circular, i.e. they do not fall on a linear scale. Instead, they are recorded on a cyclical scale, for example time of day or compass directions. Such data cannot be analyzed using standard linear statistics; therefore, circular statistical methods have been formulized to detect non-random patterns in periodically recorded data (for discussions on circular data see for example Jammalamadaka and Sengupta (2001), Mardia and Jupp (2009) and Pewsey et al. (2013)). One common example for the use of circular statistics is in orientation biology (see many examples in Batschelet (1981)). In a classic experiment, researchers would release pigeons after translocating them to a new location and test if they fly towards home (e.g. they record a vanishing bearing). To test if pigeons orient towards home, one could employ a V-test, which is by far the most powerful test in such a case where a certain heading direction (i.e. towards home) is expected (Landler et al. 2018). If the expected heading direction was unknown, a Rayleigh-test would be the most powerful alternative test of the null hypothesis of uniformity of chosen directions. A confidence interval around the mean direction can then help explore if pigeons are significantly oriented towards the home loft or deflected to a different direction.

However, in many scenarios experimenters are not only interested in testing whether a single group shows a significant direction of departure, but they want to compare animals of different origin or test the effect of experimental manipulations, e.g. to understand the underlying orientation mechanisms or cognitive abilities. Such experiments require statistical comparisons between at least two groups, e.g. one group without manipulations (the control) and one with perturbations (the experimental group). Here, we explore the robustness and power of tests designed for the comparison of two groups as the most common case, analogous to the use of a Student t-test or Mann-Whitney U-test in linear statistics.

Arguably, the most commonly-used test for differences between two circular distributions is Watson’s U^2^ test (Watson 1962), a non-parametric rank-based test that might be thought of as a circular analogue to the Mann-Whitney U-test. It is implemented in most software applications that contain circular statistic functions (e.g. Oriana (Kovach 2011), MATLAB (Berens 2009) and *R* (R Core Team 2020)). It has never been tested, however, whether the Watson U^2^ test is the most robust and powerful test across different situations. To address this, we identified 18 two-sample tests, which can test for similar null hypotheses (i.e. two circular distributions are the same). The mathematical methods underlying the different approaches vary considerably, from classical rank-sum approaches to linear statistics making use of trigonometrical functions. For brevity, we only considered test versions where the p-value is derived asymptotically rather than by randomization/bootstrapping. Our aim was to provide advice on which tests to use in which situations.

We grouped the type of comparisons between to distributions according to three possible null hypotheses: (1) The distributions are identical, (2) they have the same mean/median, or (3) they show the same concentration. Type (1) tests should be sensitive to differences in mean/median as well as to differences in the concentration, however, they should not be sensitive to differences in sample sizes. Type (2) tests should be sensitive to differences in the mean/median, but not show differences when only the concentration varies. In contrast, type (3) tests should be sensitive to differences in concentration but not detect differing means/medians.

Another level of complexity in choosing a method is that certain tests require defined prerequisites to show reliable results. The most common prerequisite is that distributions should be oriented unimodally (see for example the Watson-Williams test) or that the sample size should exceed a minimum size (e.g. this is true for the Watson’s large-sample nonparametric test) (Pewsey et al. 2013). It is unclear how robust the tests are against violation of the assumptions, but our study will offer guidance on this.

In summary, in this simulation study we explore a suite of available tests in the most common situations were two independent circular distributions are compared and subsequently derive advice for scientists working with circular data. Our analysis includes unconventional linear approaches, as well as predominantly used circular statistical tests.

## Methods

### Statistical tests used in simulation

In total we used 18 tests implemented in several *R* packages (Directional (Tsagris et al. 2020), CircStats (Lund and Agostinelli 2009), circular (Agostinelli and Lund 2017), kuiper.2samp (Ruan 2018), lawstat (Gastwirth et al. 2020), NPCirc (Oliveira Pérez et al. 2014), Rfast2 (Papadakis et al. 2019)), code for *R* functions provided in Pewsey et al. (2013), or, in one case, newly implemented for this manuscript (see table 1 and supplementary material for details on functions and packages used).

**Table 1:** List of tests used, including the functions and packages used, as well as the respective null hypothesis and test abbreviation.

| Test name | Abbreviation | Function | Package/Resource | Null hypothesis |
| --- | --- | --- | --- | --- |
| Concentration test | Con | conc.test | Directional | identical concentration |
| Embedding approach ANOVA | EmA | embed.circaov | Rfast2 | identical distribution |
| Fisher's method | FM | tang.conc | Directional | identical concentration |
| Fisher's nonparametric test | FPg | PgVal | Pewsey et al. 2013 | identical mean/median |
| Kuiper two sample test | Kui | kuiper.2samp | kuiper.2samp | identical distribution |
| Large-sample Mardia-Watson-Wheeler test | MWW | WgVal | Pewsey et al. 2013 | identical distribution |
| Levene's test | Lev | levene.test | lawstat | identical concentration |
| Log-likelihood ratio ANOVA | LIA | lr.circaov | Rfast2 | identical distribution |
| MANOVA approach | Man | manova | stats | identical distribution |
| Non equal concentration parameters approach ANOVA | HeA | het.circaov | Rfast2 | identical distribution |
| P-test | Pt | P.test.assym | Suppl in this paper | identical mean/median |
| Rao dispersion test | Rdi | rao.homogeneity | CircStats | identical concentration |
| Rao polar test | Rpo | rao.homogeneity | CircStats | identical mean/median |
| Wallraff test | Wal | wallraff.test | circular | identical concentration |
| Watson-Wheeler test | WWe | watson.wheeler.test | circular | identical distribution |
| Watson-Williams test | WWi | watson.williams.test | circular | identical mean/median |
| Watson's large-sample nonparametric test | Wat | YgVal | Pewsey et al. 2013 | identical mean/median |
| Watson's $U^2$ test | WU2 | watson.two.test | circular | identical distribution |

### Simulated distributions

All distributions were generated using the *rcircmix* function from the NPCirc package (R code in supplementary material). Each simulation was performed with 9999 randomly drawn samples from the given distribution, the proportion of significant test results is then reported as type I error or power. For all comparisons, we used the same sample size pairs per mode. For unimodal distributions we used sample sizes of 10/10, 20/20, 50/50, 20/30, 10/50, for bimodal (axial: 0° and 180°, non-symmetrical: 0° and 120°) and trimodal (symmetrical: 0°, 120° and 240°, non-symmetrical: 0°, 90° and 200°) distributions we doubled and tripled the sample sizes, respectively.

### Type I error

For each of the distributions we evaluated the type I error probability by comparing two identical distributions, with equally increasing concentration/dispersion parameters. We only continued evaluating the power of those tests that were robust, i.e. that maintained a type I error level of maximal 5% in all situations. (Note: The power of all tests was calculated and can be found in the supplementary results sheets, however, not included in the figures for clarity.)

### Power calculations – difference in concentration

In order to evaluate the power of the tests to detect differences in concentrations, we varied the concentration parameters of one of the distributions and kept the second one constant. For the von Mises distribution we used the concentration parameter κ=0 for one and values from 0 to 8 for the other distribution. For the wrapped skew normal, one distribution was kept at the dispersion parameter ρ = 1 and for the second it ranged from 1 to 4.

### Power calculations – difference in mean/median

To evaluate the power to detect differences in directionality of unimodal distributions, we kept one of the two distributions fixed at a mean of 0° and used mean directions from 0° to 180° for the second one. For bimodal distributions, we used directions from 0° to 90° and for trimodal distributions, we used 0° to 60° in order to encompass the range where we expected most differences between distributions.

### Power calculations – difference in distribution type

To test the power of the tests to differentiate between different types of distributions, we compared two distributions with similarly increasing concentration parameters across simulations (dispersion parameter in the case of the wrapped skew normal distribution), while keeping the directionality of the distributions the same between the distributions. For all the analyses, we kept the sample sizes the same as before for the respective distributions. First, we compared unimodal von Mises with an axial von Mises on the same axis. Again, the concentration parameter increased for both in the same manner. In a second simulation, we compared unimodal von Mises with unimodal wrapped skew normal distributions, using the same parameters as for the type I error calculations, for both (only the dispersion parameter range was flipped to range from 4 to 1 for the wrapped skew distribution, to follow the concentration parameter better).

### Real data examples

To test the performance of the tests in real life situations, we applied the identified robust tests to three published examples from biological studies. The first data set (*pigeon*) is included in the *R* circular package and describes a translocation experiment with homing pigeons (Gagliardo et al. 2008), where the performance of control animals was compared to pigeons with a sectioned olfactory nerve (note: a third group present in the original paper was not included in our analysis). It was expected that the control group is oriented (highly clustered) and the experimental group is disoriented (not clustered), causing a difference of concentration between the groups.

The second data set is from work by Wehner and Müller (1985) that investigated the ability of desert ants (*Cataglyphis fortis*) to transfer directional visual information received in one eye, to the other eye without further training (data included in the *R* circular package: *fisherB10c*). For the purpose of this analysis, we compared a control group (trained and tested with the same eye open) with a treatment group (trained with one eye and tested with the other eye). The hypothesis here was that information transfer between the two visual views is possible. The expectation, therefore, is a lack of difference between groups, both in direction and concentration.

The third data set is taken from a recent paper on bat navigation by Lindecke and colleagues (2019). They investigated if migratory bats (*Pipistrellus pygmaeus*) use the sun’s position at dusk to calibrate their navigational compass. For our example, we only analyzed the data from adult animals. Here the expectation was that the recorded headings should not differ in concentration, but show opposite directional preferences, according to whether they were exposed to natural or mirrored views of the sun.

## Results

### Type I error

In the case of two identical unimodal von Mises distributions with increasing concentration, several tests did not maintain type I error levels below 5% (Figure 1). The Kuiper two sample test and the “Non equal concentration parameters approach” ANOVA had increased false positive rates when the sample size was low (n=10). From the tests on identical means, the P-test, the Watson’s large-sample nonparametric test and the Watson-Williams test showed type I error rates well above 5% at larger sample sizes. From the tests on identical concentration, only the Rao dispersion test had increased type I error and only when the sample size was lower than 50. The type I error results were similar for the unimodal wrapped skew normal distribution, however with the Wallraff test and Fisher’s method showing additional type I error inflation (Figure S1).

**Figure 1:**
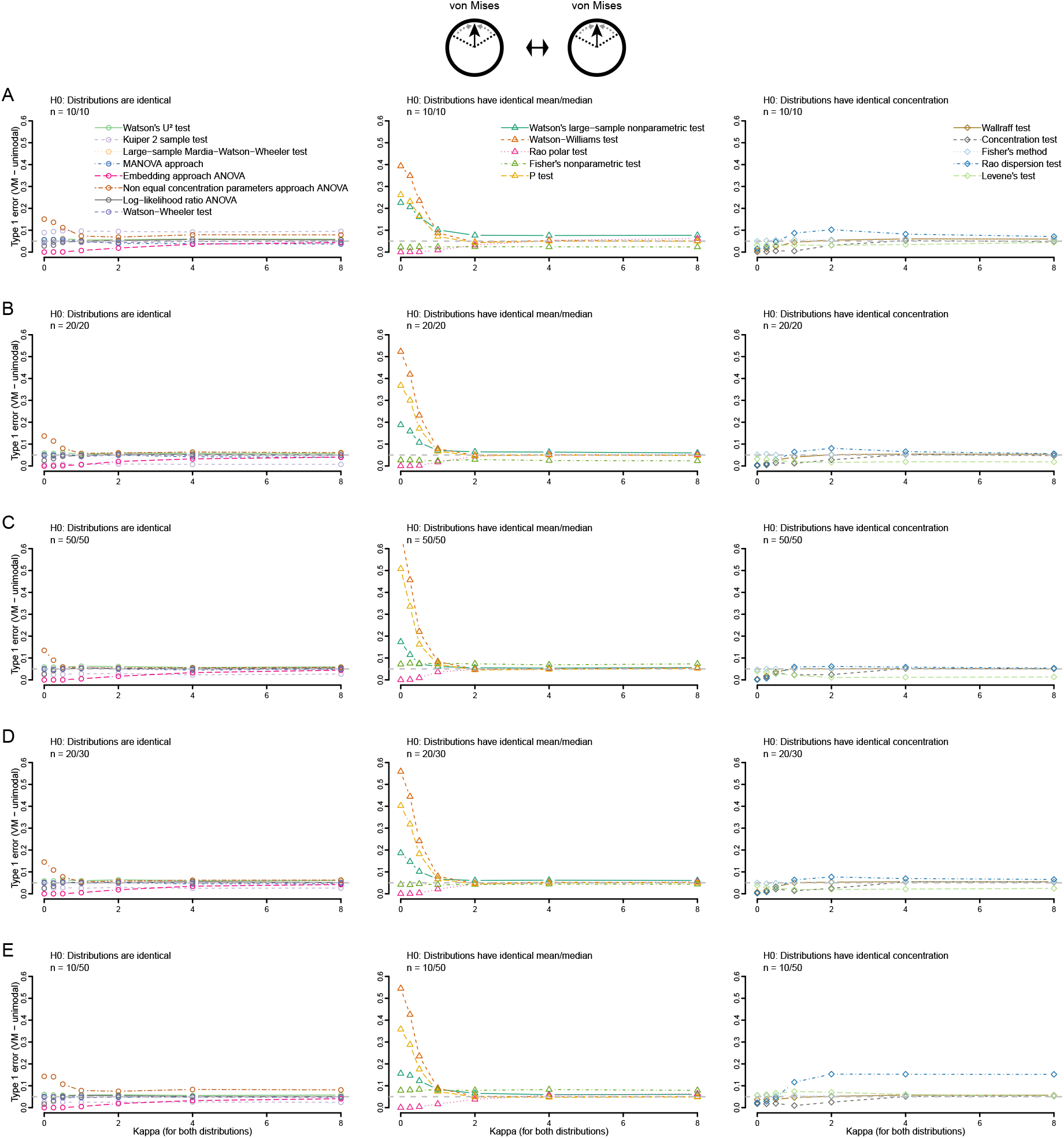
Type 1 error of all tests using von Mises distributions for different sample sizes: 10 and 10 (A), 20 and 20 (B), 50 and 50 (C), 20 and 30 (D) and 10 and 50 (E). Concentration (κ, kappa) increases for both distributions from 0 to 8. Tests are grouped according to their null hypotheses.

When we tested two identical axial distributions, five tests failed to maintain 5% type I error levels: the P-test, the Watsons large sample nonparametric test, the Fisher’s method, the Watson-Williams test and the “Non equal concentration parameters approach” ANOVA (Figure S2). In the case of two identical asymmetrical bimodal distributions seven tests failed to maintain the expected type I error rates: the P-test, the Watsons large sample nonparametric test, the Fisher’s method, the Watson-Williams test, the “Non equal concentration parameters approach” ANOVA, the Wallraff test and the Log-likelihood ratio ANOVA (Figure S3). For two trimodal situations (symmetrical (Figure S4) and asymmetrical (Figure S5)), the results were comparable to the bimodal distributions, but now Fisher’s nonparametric test also showed slightly increased type I error rates. One of the main issues for most tests appeared to be related to the symmetric components of the multimodal distributions.

In sum, only eight out of 18 tests were robust, i.e. they showed no issues regarding increased type I error probabilities across the situations investigated: the Watson U^2^ test, the Rao polar test, the Large-sample Mardia-Watson-Wheeler test, the concentration test, the Watson-Wheeler test, the embedding approach ANOVA, the MANOVA approach and the Levene’s test. We continued to explore the power of these eight robust tests.

### Power to detect differences in concentration

The most powerful test to detect concentration differences between two von Mises distributions was the MANOVA approach, which offered superior power especially at lower sample sizes (Figure 2). The Watson U^2^ test was also very powerful, followed by the Watson-Wheeler and the Large-sample Mardia-Watson-Wheeler tests with only marginally lower power. The embedding approach ANOVA had lower power, but, notably, was still more powerful than the Concentration test and Levene’s test, both specifically designed to detect differences in concentration. As expected, the Rao polar test was not sensitive to differences in concentration. The general results for two unimodal wrapped skew normal distributions were comparable to the results for unimodal von Mises distributions, with the only exception of superior performance of Levene’s test in situations with highly asymmetric samples sizes (Figure S6).

**Figure 2:**
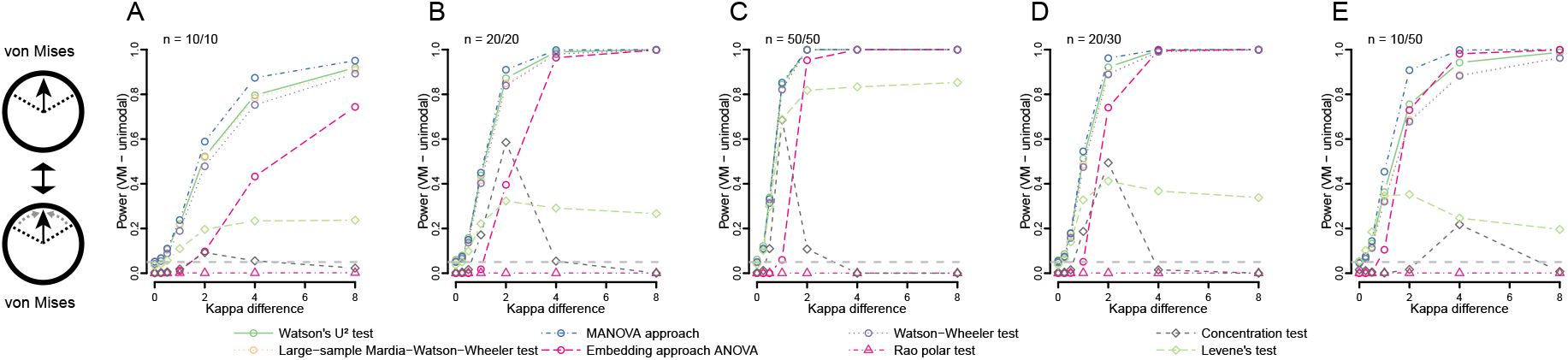
Power of all included tests when comparing von Mises distributions of differing concentrations using different sample sizes: 10 and 10 (A), 20 and 20 (B), 50 and 50 (C), 20 and 30 (D) and 10 and 50 (E). The first distribution is fixed at κ = 0, the second increases from 0 towards 8.

When comparing axial von Mises distributions, only the Watson U^2^ test offered acceptable power (Figure S7). For the symmetrical trimodal distributions, overall power was very low, and again, only the Watson U^2^ providing some power (Figure S8). The asymmetrical bimodal (Figure S9) situation showed acceptable power of the MANOVA approach and Watson’s U^2^, however, for the asymmetrical trimodal power was low with the Watson’s U^2^ providing the best results (Figure S10).

### Power to detect differences in the mean/median

The power to detect angular differences between two von Mises distributions was highest for the MANOVA approach at small sample sizes (n=10), followed by the Watson U^2^ and Watson-Wheeler test and the Large-sample Mardia-Watson-Wheeler test (Figure 3). Interestingly, the Levene’s test also showed lower but acceptable power levels, although it was expected to be sensitive to concentration differences (to which it was less sensitive, see Figure 2). The concentration test was not sensitive to the differences in mean direction. Special cases were the embedding approach ANOVA and the Rao polar test. The ANOVA approach showed, with the exception of very unequal sample sizes (n=10/50), a unimodal response, with increasing power levels from 0° to 90° difference, but then rapidly decreasing power towards 180° difference. The Rao polar test showed an even stranger pattern, with, at higher sample sizes, very good power when the difference was either around 45° or 135°, but with power levels dropping to 0.05 in between these two peaks (at 90°). The results were similar for the wrapped skew normal distribution, with the exceptions that the Rao polar test showed strongly reduced power and switched from a bimodal to a unimodal power curve with a peak around 60°, and the Levene’s test completely lost its power (Figure S11).

**Figure 3:**
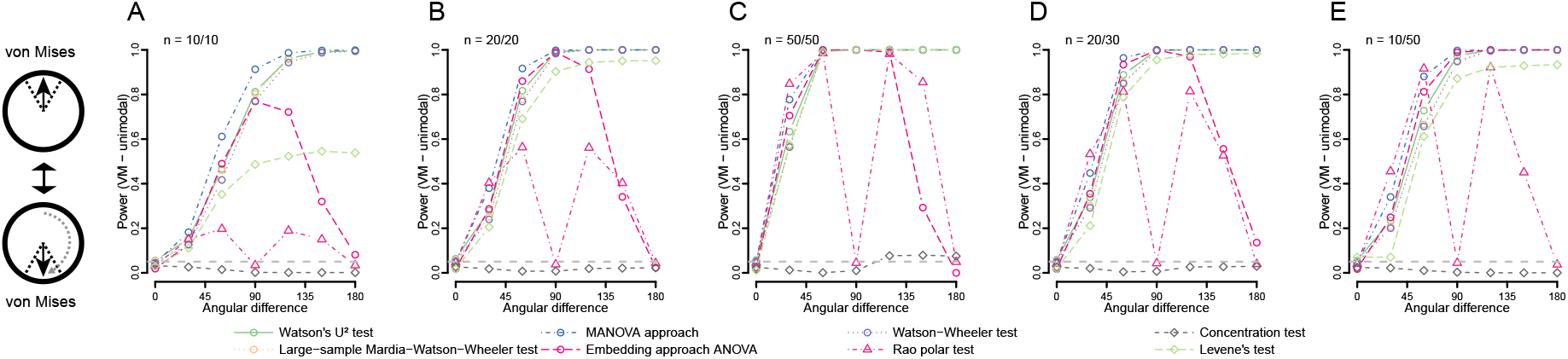
Power of all included tests when comparing von Mises distributions (kappa for both = 2) of differing directions using different sample sizes: 10 and 10 (A), 20 and 20 (B), 50 and 50 (C), 20 and 30 (D) and 10 and 50 (E). The first distribution is fixed at 0°, the second increases from 0° towards 180.

For axial distributions, only the Watson U^2^ test offered acceptable power levels, although large sample sizes (~n=100) were required for the power to reach over 50% (Figure S12). All other tests failed to detect the difference in mean direction between two axial distributions. For symmetric trimodal distributions none of the tests used was sensitive to differences in mean direction (Figure S13).

When comparing asymmetrical bimodal distributions, the general trends were similar to the unimodal case. However, over all sample sizes the MANOVA approach offered the best power. The Watson-Wheeler test was considerably less powerful in this situation, as were the Watson U^2^ test and the Large-sample Mardia-Watson-Wheeler test (Figure S14). The Levene’s test showed a unimodal -shaped power curve. The asymmetrical trimodal situation was, again, similar to the asymmetrical bimodal situation (Figure S15), with the exception of the Levene’s test, which showed steady power increase with angular difference (instead of the hump-shaped curve).

### Power to detect differences in distribution type

When comparing a unimodal and an axial bimodal distribution, which increased similarly in concentration, we found that the MANOVA approach again offered the best power in particular at low samples sizes, followed by the Watson U^2^ test, the Large-sample Mardia-Watson-Wheeler test and Watson-Wheeler test (Figure 4). While the embedding approach ANOVA and the Levene’s test had varying but usable power levels, the concentration test was only sensitive to such differences at low concentration values. The Rao polar test was not sensitive to such differences.

**Figure 4:**
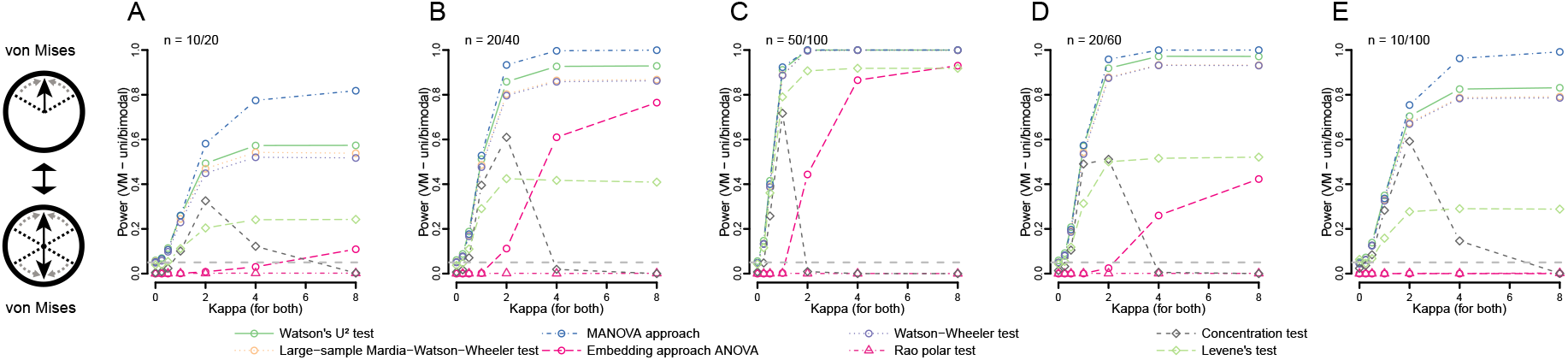
Power of all included tests when comparing von Mises distributions of differing number of modes (one and axially bimodal) using different sample sizes: 10 and 20 (A), 20 and 40 (B), 50 and 100 (C), 20 and 60 (D) and 10 and 100 (E). The concentration (κ) of both increases from 0 to 8.

The picture was only marginally different when comparing a von Mises with a wrapped skew normal distribution (Figure S16). For low sample sizes (n=10) the MANOVA approach offered great power, followed by the embedding approach ANOVA. The latter offered good power throughout the range of sample sizes tested, followed by the Watson U^2^ test, the Large-sample Mardia-Watson-Wheeler test and Levene’s test. Also, the Rao polar test showed lower, but acceptable sensitivity to distribution type. The concentration test only showed very low power, that (as expected) increased with increasing concentrations of the respective distributions.

### Real data examples

Testing the performance of the robust tests on real data sets revealed, predominantly, the expected test behavior. In the example of homing pigeons where a difference in concentration was expected, all tests, with the exception of the Rao polar test and, notably, the concentration test, showed a significant difference between the distributions (Figure 5A). Therefore, we can conclude, in accordance with the respective publication (Gagliardo et al. 2008), that sectioning of the olfactory nerve disrupted the homing behavior of pigeons.

**Figure 5:**
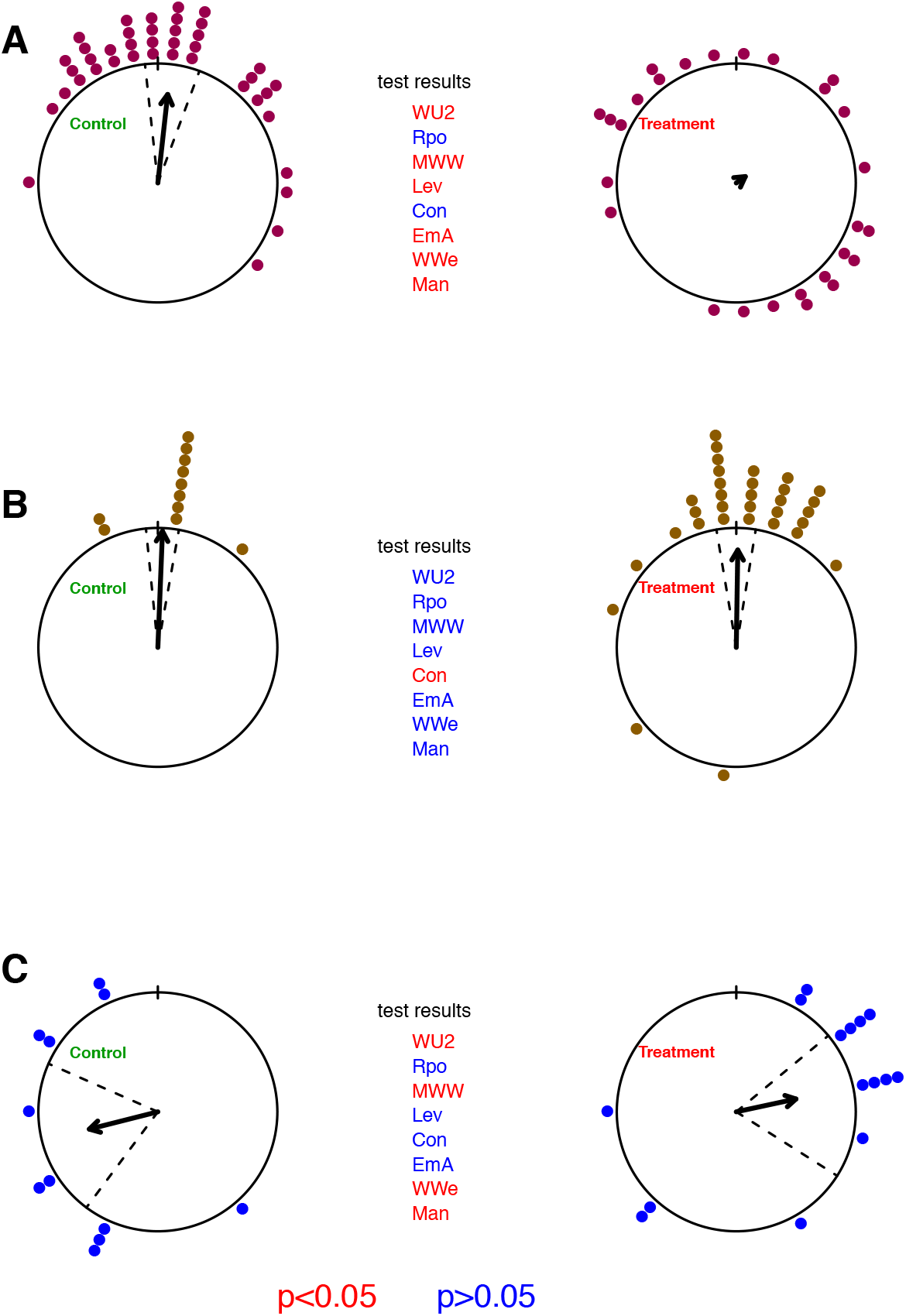
Results from example data. Shown are results of pigeon (A), ant (B) and bat orientations (C). Control groups are on the left panels and experimental groups on the right. The tests are abbreviated according to table 1, significant test results are indicated in red and non-significant in blue. For each circular plot directional data is shown as dots on the circle (each dot is one individual), the arrows represent the mean direction and the dashed line the 95% confidence interval.

In the ant example, where no difference between the groups was expected, there was no significant difference between the distributions detected by most of the tests (Figure 5B). Only the concentration test showed a significant difference. Based on the other tests we would conclude that there was no biological meaningful difference between the two distributions. Therefore, ants appear to be able to transfer visual information from one eye to the other.

In the bat example, where a difference in mean direction was expected, the Watson U^2^, the Mardia-Watson-Wheeler, Watson-Wheeler test and the MANOVA approach showed a significant difference (Figure 5C). Notably, the Rao polar, Levene’s, and concentration tests and the embedding approach ANOVA failed to show a significant difference. At least for the Rao polar test, one would have expected a significant difference, as the two distributions are clearly 180° apart. This outcome concurs with our simulation results where the Rao polar test failed to distinguish distributions on the same axis (Figure 8). As the results of the tests where quite mixed this example highlights the need for choosing a test with appropriate power to detect the expected differences. Based on the results of the most powerful tests, we conclude that the bats showed a mirrored orientation, as expected in the experimental design.

## Discussion

Our extensive power analyses of tests for analyzing if two circular distributions are the same show that many of the tests suffer from elevated type I error rates, at least in specific situations. To minimize the risk of getting false positive results we recommend not using any of these tests. Of the eight remaining tests, only one test, the Watson U^2^ test, offers good power in all of the tested situations. An interesting alternative revealed by our analysis, however, is the MANOVA approach, which is also very powerful, and furthermore allows the addition of covariates.

To our knowledge, the MANOVA approach, as used in this analysis, has only been used once in similar fashion in order to test the effect of environmental factors on toad orientation (Pail et al. 2020). Nobody seems to have used it for distribution comparisons. This is surprising, as the underlying ideas (transforming an angle in two linear components using trigonometric functions) and the MANOVA calculation itself are not at all new to statistics. Given the potential advantages of such an approach in terms of adding covariates and its robust and powerful behavior we would recommend the broader uptake of this test in circular statistics when it comes to group comparisons.

Some of the tests that we did not evaluate further might be very powerful for a certain set of situations, where they maintain expected type I error rates. However, such utilization of statistical tests can easily lead to false positive results and in general might be quite delicate to handle for the experimenter. They would have to inspect their distributions closely before testing. Such test selection on the basis of visual inspection of the data by itself could lead to increased type I error rates. In contrast to linear stats where certain distributions can be expected by study design (e.g. random sampling of continuous variable would suggest a Gaussian distribution, or count data a Poisson distribution), this is not the case for circular statistics.

The behavior of the concentration test is puzzling, in general it has low power to detect differences, but appears to be highly sensitive to very specific differences. From inspection of the results of the ant example (see Figure 5B) it is clear that the two distributions are not identical in terms of clustering, although the confidence intervals and mean vector lengths are similar. In the control group most animals are exactly at the same direction, while there is a more even spread in the experimental distribution. Therefore, the detected difference might be real, but it is unclear what feature of the data exactly this test detects.

In case of expected axial orientation for both distribution we recommend doubling the angles and reducing to modulo 360° for both distributions prior to applying any of the tests, in order to overcome the extreme loss of statistical power in such situations (e.g. Figure S7). If this is not possible or attractive to the researcher, the best option is using Watson’s U^2^ test.

In the current study, we only tested two distributions against each other, although six out of the eight tests included in the power analysis would also allow testing of multiple distributions. We surmise that it is safe to assume that the general power comparisons between those tests would show similar patterns also when considering more groups. Exploring the performance of tests using multiple treatment groups would be a fruitful avenue of further research, as would evaluation of paired data.

## Supporting information

Supplementary material, code and results

## Acknowledgements

LL is supported by the Austrian Science Fund (FWF, Grant Number: P32586).

## Authors’ contributions

LL, GDR and EPM conceptualized the problem and discussed the analytic approaches. LL and GDR prepared the code. LL, GDR and EPM interpreted the results. GDR, LL and EPM wrote the manuscript.

## Availability of data and materials

Data and code are available in the supplementary material.

