## Supplementary material, code and results for "Comparing two circular distributions: advice for effective implementation of statistical procedures in biology": Biological_Examples_TwoSamples_OnlySubset_of_tests_PascalEdit.R

```
library(kuiper.2samp)
library(lawstat)
library(circular)
library(compiler)
library(Directional)
library(NPCirc)
library(CircStats)
library(Rfast2)

source("Functions_JRSB_March2021.R")

#######################################
#####Data sets 
#######################################


#Gagliardo A., Ioale' P., Savini M., and Wild M. (2008). 
#avigational abilities of homing pigeons deprived of olfactory or trigeminally mediated magnetic information when young. 
#J. Exp. Biol., 211:2046–2051.

data(pigeons) # in package circular 
pigeons$bearing<-rad(pigeons$bearing)
pigeons_c<-pigeons$bearing[pigeons$treatment=="c"]
pigeons_on<-pigeons$bearing[pigeons$treatment=="on"]

#R. Wehner and M. Muller (1985) 
#Does interocular transfer occur in visual navigation by ants? 
#Nature, 315, 228-9.

data(fisherB10c)
Ants_1 <- rad(fisherB10c[["set1"]]) #Fig1A
Ants_2 <- rad(fisherB10c[["set2"]]) #Fig1B1

#Lindecke et al. Current biology
bats<-read.csv("rawdata_lindecke.csv")

bats<-bats[bats$age=="adult",]#only take adults
bats$takeoff.orientation<-rad(bats$takeoff.orientation)
bats_C<-bats$takeoff.orientation[bats$treatment=="C"]
bats_M<-bats$takeoff.orientation[bats$treatment=="M"]


#######################################
#####P-value calculation
#######################################

Distribution1_list = list(pigeons_c,Ants_1,bats_C)
Distribution2_list = list(pigeons_on,Ants_2,bats_M)
  
#preparing results file 
Results_Examples <- data.frame(matrix(nrow = length(Distribution1_list), ncol = 0))

#starting the loop over all the data sets 

for (i in 1:length(Distribution1_list)){

Distribution1<-Distribution1_list[[i]]
Distribution2<-Distribution2_list[[i]]

nv1<-length(Distribution1)
nv2<-length(Distribution2)

#Making treatment column
Group1 <-(as.matrix( rep((as.matrix((strrep(c("A"), 1)))),nv1),dim=c(nv1, 1)))
Group2 <- (as.matrix( rep((as.matrix((strrep(c("B"), 1)))),nv2),dim=c(nv2, 1)))
GroupsAB<-as.data.frame(c(Group1[,1], Group2[,1]))

##watson2SampleTest
watson1 = watson.two.test(Distribution1,Distribution2)
watson1 <- array(as.matrix(unlist(watson1)), dim=c(5, 1))
Results_Examples$WU2[i]<- if (watson1[1,]>0.178) (0.04) else (0.06)  # Critical Value: 0.187 #this is just to have number higher and one lower than 0.05 (see coding below)

#Raos Tests for Homogeneity
rao1 = capture.output(rao.homogeneity(list(Distribution1, Distribution2),alpha=0.05))
Results_Examples$Rpo[i]<- if (rao1[9] == "Reject null hypothesis of equal dispersions ") (0.04) else (0.06) #this is just to have number higher and one lower than 0.05 (see coding below)

#####preparing sample in the way I need it for the tests in the Pewsey et al. - and getting ndat and g
DistGroups<-c(Distribution1, Distribution2)
ndat <- c(nv1, nv2)
g <- length(ndat)


# Large-sample Mardia-Watson-Wheeler test for a common distribution
Results_Examples$MWW[i]<- pchisq(WgVal(CosSinUniScores(DistGroups), ndat, g), 2*(g-1), lower.tail=F)

#Levenes # 
LevenesRES <-  lawstat::levene.test(DistGroups,GroupsAB[,1])
Results_Examples$Lev[i] <- LevenesRES["p.value"]

#FromDirectionalPackage  Test of equality of the concentration parameters for circular data from Mardia, K. V. and Jupp, P. E. (2000). Directional statistics. Chicester: John Wiley & Sons.
conctestDir <- conc.test(DistGroups,as.factor(GroupsAB[,1]),rads=TRUE)
Results_Examples$Con[i]<- conctestDir$res["p-value"]

#embed.circaov from Rfast2 (originally from Directional)
embed.circaovRfast2 <-  Rfast2::embed.circaov(DistGroups,as.factor(GroupsAB[,1]))
Results_Examples$EmA[i]<-embed.circaovRfast2[2]

#Watson wheeler test! Test assumes von Mises
watson.wheelerRES <-  watson.wheeler.test(DistGroups,GroupsAB[,1])
Results_Examples$WWe[i] <- watson.wheelerRES["p.value"]

##MANOVA approach 
ManCirc<-summary(manova(cbind(cos(DistGroups),sin(DistGroups)) ~ GroupsAB[,1]))
Results_Examples$Man[i] <- ManCirc$stats[1,6]

}#i

#prepare color and names for insert in figure 
Testabbreviations<-colnames(Results_Examples)
colvector_pigeons<- sapply(Results_Examples[1,],FUN=function(x) (if (x<0.05) ("red") else ("blue"))) 
colvector_ants<- sapply(Results_Examples[2,],FUN=function(x) (if (x<0.05) ("red") else ("blue"))) 
colvector_bats<- sapply(Results_Examples[3,],FUN=function(x) (if (x<0.05) ("red") else ("blue"))) 

######################################################################
####Plot
######################################################################

# library(extrafont) # in order to use Arial for figures 
# font_import() # run this line if fonts are not yet imported 
# loadfonts(device="pdf") 

library(shape) # in order to use different kind of arrows for the plots 

#load my own functions
source("Functions_JRSB_March2021.R")

#predfine colors for all plots 
H ="#FF8000";control = "#009900";trt="red" ; pcolor = "darkorchid3";pigeons_col = "#99004C"
ants_col ="orange4" ; bats_col = "blue"  ;D = "#0000FF";cexp = 0.8

pdf("Figure 5.pdf", width=4.98, height=6)
par(mar=c(0,0,0,0),mfrow = c(3, 2))

  #define circular plotting parameters
  sep=0.1 #distance between dots
  shrink=1.8 # size parameter for circle 
  cexD = 1 # general size parameter for basic plot.circular 
  start.sep = 0.08 #distance between dots and circle 
  cexspecies=0.65 #size of text placed inside the circle "Control" / "Trt"
  cexlabel=0.6 #label size ("N" or "0" or such)
  tcltext=0.1 #spacing between tick and label 
  lwdarrow=2 #arrow width
  lengtharrow=0.05 #arrow length
  tclength=0.1 # tick length
  
  textTestX=1.5;textTestY=1
  
  # A ####### Pigeon Data - Control
  
  wrapper_unimodal_plot(pigeons_c,pigeons_col,labels=c(""))
  
  text(-1.2,1.2,"A",font=2,cex=1.5) 
  
  text(-0.4,.5,"Control",cex = cexspecies,col=control,font=2)

  
  legend(x=textTestX+0.3, y=textTestY,Testabbreviations,bty="n",
         text.col=colvector_pigeons,cex=0.8 ,xpd=NA,ncol=1,
         title="test results",title.col = "black")
  # B ####### Pigeon Data - ON
  
  wrapper_unimodal_plot(pigeons_on,pigeons_col,labels=c(""))
 
   text(-0.4,.5,"Treatment",cex = cexspecies,col=trt,font=2)
  
  
  # C ####### Ant Data - Control
  
  wrapper_unimodal_plot(Ants_1,ants_col,labels=c(""))
  
  text(-1.2,1.2,"B",font=2,cex=1.5) 
  
  text(-0.4,.5,"Control",cex = cexspecies,col=control,font=2)
 
   legend(x=textTestX+0.3, y=textTestY,Testabbreviations,bty="n",
         text.col=colvector_ants,cex=0.8 ,xpd=NA,ncol=1,
         title="test results",title.col = "black")
  
  # D ####### Ant Data - Treatment
  
  wrapper_unimodal_plot(Ants_2,ants_col,labels=c(""))
 
    text(-0.45,.5,"Treatment",cex = cexspecies,col=trt,font=2)
  
  # E ####### Bat Data - Control
  
  wrapper_unimodal_plot(bats_C,bats_col,labels=c(""))
 
   legend(x=textTestX-1, y=textTestY-2.2,c("p<0.05","p>0.05"),bty="n",
         text.col=c("red", "blue"),cex=1.3 ,xpd=NA,ncol=2,
  )
   
  text(-1.2,1.2,"C",font=2,cex=1.5) 
  text(-0.4,.5,"Control",cex = cexspecies,col=control,font=2)
  legend(x=textTestX+0.3, y=textTestY,Testabbreviations,bty="n",
         text.col=colvector_bats,cex=0.8 ,xpd=NA,ncol=1,
         title="test results",title.col = "black")
# E ####### Bat Data - M

wrapper_unimodal_plot(bats_M,bats_col,labels=c(""))

text(-0.4,.5,"Treatment",cex = cexspecies,col=trt,font=2)


dev.off()
```
