## Supplementary material, code and results for "Comparing two circular distributions: advice for effective implementation of statistical procedures in biology": FigureAll_Power_plotting_feb2021_Pascaledit_WSNnormal.R

```
#Plotting Comp Dist

#Figure 1 (VM - identical distributions): A: (10,10), B: (20,20), C: (50,50),D: (20,30), E: (10,50), 

#load data 
Figure1PA=read.csv("Results/Power_n1_10_n2_10_its_9999_Dist1_vm_Dist2_vm_m1=0-0_m2_0-0_k1=0-0_k2_0-8.csv")
Figure1PB=read.csv("Results/Power_n1_20_n2_20_its_9999_Dist1_vm_Dist2_vm_m1=0-0_m2_0-0_k1=0-0_k2_0-8.csv")
Figure1PC=read.csv("Results/Power_n1_50_n2_50_its_9999_Dist1_vm_Dist2_vm_m1=0-0_m2_0-0_k1=0-0_k2_0-8.csv")
Figure1PD=read.csv("Results/Power_n1_20_n2_30_its_9999_Dist1_vm_Dist2_vm_m1=0-0_m2_0-0_k1=0-0_k2_0-8.csv")
Figure1PE=read.csv("Results/Power_n1_10_n2_50_its_9999_Dist1_vm_Dist2_vm_m1=0-0_m2_0-0_k1=0-0_k2_0-8.csv")

list_fig1 <- list(Figure1PA, Figure1PB,Figure1PC,Figure1PD,Figure1PE)

Figure2PA=read.csv("Results/Power_n1_10_n2_10_its_9999_Dist1_vm_Dist2_vm_m1=0-0_m2_0-180_k1=2-2_k2_2-2.csv")
Figure2PB=read.csv("Results/Power_n1_20_n2_20_its_9999_Dist1_vm_Dist2_vm_m1=0-0_m2_0-180_k1=2-2_k2_2-2.csv")
Figure2PC=read.csv("Results/Power_n1_50_n2_50_its_9999_Dist1_vm_Dist2_vm_m1=0-0_m2_0-180_k1=2-2_k2_2-2.csv")
Figure2PD=read.csv("Results/Power_n1_20_n2_30_its_9999_Dist1_vm_Dist2_vm_m1=0-0_m2_0-180_k1=2-2_k2_2-2.csv")
Figure2PE=read.csv("Results/Power_n1_10_n2_50_its_9999_Dist1_vm_Dist2_vm_m1=0-0_m2_0-180_k1=2-2_k2_2-2.csv")

list_fig2 <- list(Figure2PA, Figure2PB,Figure2PC,Figure2PD,Figure2PE)

# 
Figure3PA=read.csv("Results/Power_vm_vmaxial_n1_10_n2_20_its_9999_Dist1_vm_Dist2_vm_m1=0-0_m2_0-0_k1=0-8_k2_0-8.csv")
Figure3PB=read.csv("Results/Power_vm_vmaxial_n1_20_n2_40_its_9999_Dist1_vm_Dist2_vm_m1=0-0_m2_0-0_k1=0-8_k2_0-8.csv")
Figure3PC=read.csv("Results/Power_vm_vmaxial_n1_50_n2_100_its_9999_Dist1_vm_Dist2_vm_m1=0-0_m2_0-0_k1=0-8_k2_0-8.csv")
Figure3PD=read.csv("Results/Power_vm_vmaxial_n1_20_n2_60_its_9999_Dist1_vm_Dist2_vm_m1=0-0_m2_0-0_k1=0-8_k2_0-8.csv")
Figure3PE=read.csv( "Results/Power_vm_vmaxial_n1_10_n2_100_its_9999_Dist1_vm_Dist2_vm_m1=0-0_m2_0-0_k1=0-8_k2_0-8.csv")
# 
 list_fig3 <- list(Figure3PA, Figure3PB,Figure3PC,Figure3PD,Figure3PE)
# 
Figure4PA=read.csv("Results/Power_vm_wsn_n1_10_n2_10_its_9999_Dist1_vm_Dist2_wsn_m1=0-0_m2_0-0_k1=0-8_k2_1-4.csv")
Figure4PA$x=Figure1PA$x
Figure4PB=read.csv("Results/Power_vm_wsn_n1_20_n2_20_its_9999_Dist1_vm_Dist2_wsn_m1=0-0_m2_0-0_k1=0-8_k2_1-4.csv")
Figure4PB$x=Figure1PA$x
Figure4PC=read.csv("Results/Power_vm_wsn_n1_50_n2_50_its_9999_Dist1_vm_Dist2_wsn_m1=0-0_m2_0-0_k1=0-8_k2_1-4.csv")
Figure4PC$x=Figure1PA$x
Figure4PD=read.csv("Results/Power_vm_wsn_n1_20_n2_30_its_9999_Dist1_vm_Dist2_wsn_m1=0-0_m2_0-0_k1=0-8_k2_1-4.csv")
Figure4PD$x=Figure1PA$x
Figure4PE=read.csv( "Results/Power_vm_wsn_n1_10_n2_50_its_9999_Dist1_vm_Dist2_wsn_m1=0-0_m2_0-0_k1=0-8_k2_1-4.csv")
Figure4PE$x=Figure1PA$x
# 
list_fig4 <- list(Figure4PA, Figure4PB,Figure4PC,Figure4PD,Figure4PE)

#load data 
Figure5PA=read.csv("Results/Power_n1_10_n2_10_its_9999_Dist1_wsn_Dist2_wsn_m1=0-0_m2_0-0_k1=1-1_k2_1-4.csv")
Figure5PB=read.csv("Results/Power_n1_20_n2_20_its_9999_Dist1_wsn_Dist2_wsn_m1=0-0_m2_0-0_k1=1-1_k2_1-4.csv")
Figure5PC=read.csv("Results/Power_n1_50_n2_50_its_9999_Dist1_wsn_Dist2_wsn_m1=0-0_m2_0-0_k1=1-1_k2_1-4.csv")
Figure5PD=read.csv("Results/Power_n1_20_n2_30_its_9999_Dist1_wsn_Dist2_wsn_m1=0-0_m2_0-0_k1=1-1_k2_1-4.csv")
Figure5PE=read.csv("Results/Power_n1_10_n2_50_its_9999_Dist1_wsn_Dist2_wsn_m1=0-0_m2_0-0_k1=1-1_k2_1-4.csv")

list_fig5 <- list(Figure5PA, Figure5PB,Figure5PC,Figure5PD,Figure5PE)

Figure6PA=read.csv("Results/Power_n1_10_n2_10_its_9999_Dist1_wsn_Dist2_wsn_m1=0-0_m2_0-180_k1=2-2_k2_2-2.csv")
Figure6PB=read.csv("Results/Power_n1_20_n2_20_its_9999_Dist1_wsn_Dist2_wsn_m1=0-0_m2_0-180_k1=2-2_k2_2-2.csv")
Figure6PC=read.csv("Results/Power_n1_50_n2_50_its_9999_Dist1_wsn_Dist2_wsn_m1=0-0_m2_0-180_k1=2-2_k2_2-2.csv")
Figure6PD=read.csv("Results/Power_n1_20_n2_30_its_9999_Dist1_wsn_Dist2_wsn_m1=0-0_m2_0-180_k1=2-2_k2_2-2.csv")
Figure6PE=read.csv("Results/Power_n1_10_n2_50_its_9999_Dist1_wsn_Dist2_wsn_m1=0-0_m2_0-180_k1=2-2_k2_2-2.csv")

list_fig6 <- list(Figure6PA, Figure6PB,Figure6PC,Figure6PD,Figure6PE)

Figure7PA=read.csv("Results/Power_n1_20_n2_20_its_9999_Dist1_Axialvm_Dist2_vm_m1=0-0_m2_0-0_k1=0-0_k2_0-8.csv")
Figure7PB=read.csv("Results/Power_n1_40_n2_40_its_9999_Dist1_Axialvm_Dist2_vm_m1=0-0_m2_0-0_k1=0-0_k2_0-8.csv")
Figure7PC=read.csv("Results/Power_n1_100_n2_100_its_9999_Dist1_Axialvm_Dist2_vm_m1=0-0_m2_0-0_k1=0-0_k2_0-8.csv")
Figure7PD=read.csv("Results/Power_n1_40_n2_60_its_9999_Dist1_Axialvm_Dist2_vm_m1=0-0_m2_0-0_k1=0-0_k2_0-8.csv")
Figure7PE=read.csv( "Results/Power_n1_20_n2_100_its_9999_Dist1_Axialvm_Dist2_vm_m1=0-0_m2_0-0_k1=0-0_k2_0-8.csv")

list_fig7 <- list(Figure7PA, Figure7PB,Figure7PC,Figure7PD,Figure7PE)

Figure8PA=read.csv("Results/Power_n1_20_n2_20_its_9999_Dist1_Axialvm_Dist2_vm_m1=0-0_m2_0-90_k1=2-2_k2_2-2.csv")
Figure8PB=read.csv("Results/Power_n1_40_n2_40_its_9999_Dist1_Axialvm_Dist2_vm_m1=0-0_m2_0-90_k1=2-2_k2_2-2.csv")
Figure8PC=read.csv("Results/Power_n1_100_n2_100_its_9999_Dist1_Axialvm_Dist2_vm_m1=0-0_m2_0-90_k1=2-2_k2_2-2.csv")
Figure8PD=read.csv("Results/Power_n1_40_n2_60_its_9999_Dist1_Axialvm_Dist2_vm_m1=0-0_m2_0-90_k1=2-2_k2_2-2.csv")
Figure8PE=read.csv( "Results/Power_n1_20_n2_100_its_9999_Dist1_Axialvm_Dist2_vm_m1=0-0_m2_0-90_k1=2-2_k2_2-2.csv")

list_fig8 <- list(Figure8PA, Figure8PB,Figure8PC,Figure8PD,Figure8PE)

Figure9PA=read.csv("Results/Power_n1_30_n2_30_its_9999_Dist1_Symm_trimodalvm_Dist2_vm_m1=0-0_m2_0-0_k1=0-0_k2_0-8.csv")
Figure9PB=read.csv("Results/Power_n1_60_n2_60_its_9999_Dist1_Symm_trimodalvm_Dist2_vm_m1=0-0_m2_0-0_k1=0-0_k2_0-8.csv")
Figure9PC=read.csv("Results/Power_n1_150_n2_150_its_9999_Dist1_Symm_trimodalvm_Dist2_vm_m1=0-0_m2_0-0_k1=0-0_k2_0-8.csv")
Figure9PD=read.csv("Results/Power_n1_60_n2_90_its_9999_Dist1_Symm_trimodalvm_Dist2_vm_m1=0-0_m2_0-0_k1=0-0_k2_0-8.csv")
Figure9PE=read.csv( "Results/Power_n1_30_n2_150_its_9999_Dist1_Symm_trimodalvm_Dist2_vm_m1=0-0_m2_0-0_k1=0-0_k2_0-8.csv")

list_fig9 <- list(Figure9PA, Figure9PB,Figure9PC,Figure9PD,Figure9PE)

Figure10PA=read.csv("Results/Power_n1_30_n2_30_its_9999_Dist1_Symm_trimodalvm_Dist2_vm_m1=0-0_m2_0-60_k1=2-2_k2_2-2.csv")
Figure10PB=read.csv("Results/Power_n1_60_n2_60_its_9999_Dist1_Symm_trimodalvm_Dist2_vm_m1=0-0_m2_0-60_k1=2-2_k2_2-2.csv")
Figure10PC=read.csv("Results/Power_n1_150_n2_150_its_9999_Dist1_Symm_trimodalvm_Dist2_vm_m1=0-0_m2_0-60_k1=2-2_k2_2-2.csv")
Figure10PD=read.csv("Results/Power_n1_60_n2_90_its_9999_Dist1_Symm_trimodalvm_Dist2_vm_m1=0-0_m2_0-60_k1=2-2_k2_2-2.csv")
Figure10PE=read.csv( "Results/Power_n1_30_n2_150_its_9999_Dist1_Symm_trimodalvm_Dist2_vm_m1=0-0_m2_0-60_k1=2-2_k2_2-2.csv")

list_fig10 <- list(Figure10PA, Figure10PB,Figure10PC,Figure10PD,Figure10PE)

Figure11PA=read.csv("Results/Power_n1_20_n2_20_its_9999_Dist1_Asymm_bimodalvm_Dist2_vm_m1=0-0_m2_0-0_k1=0-0_k2_0-8.csv")
Figure11PB=read.csv("Results/Power_n1_40_n2_40_its_9999_Dist1_Asymm_bimodalvm_Dist2_vm_m1=0-0_m2_0-0_k1=0-0_k2_0-8.csv")
Figure11PC=read.csv("Results/Power_n1_100_n2_100_its_9999_Dist1_Asymm_bimodalvm_Dist2_vm_m1=0-0_m2_0-0_k1=0-0_k2_0-8.csv")
Figure11PD=read.csv("Results/Power_n1_40_n2_60_its_9999_Dist1_Asymm_bimodalvm_Dist2_vm_m1=0-0_m2_0-0_k1=0-0_k2_0-8.csv")
Figure11PE=read.csv( "Results/Power_n1_20_n2_100_its_9999_Dist1_Asymm_bimodalvm_Dist2_vm_m1=0-0_m2_0-0_k1=0-0_k2_0-8.csv")

list_fig11 <- list(Figure11PA, Figure11PB,Figure11PC,Figure11PD,Figure11PE)

Figure12PA=read.csv("Results/Power_n1_20_n2_20_its_9999_Dist1_Asymm_bimodalvm_Dist2_vm_m1=0-0_m2_0-90_k1=2-2_k2_2-2.csv")
Figure12PB=read.csv("Results/Power_n1_40_n2_40_its_9999_Dist1_Asymm_bimodalvm_Dist2_vm_m1=0-0_m2_0-90_k1=2-2_k2_2-2.csv")
Figure12PC=read.csv("Results/Power_n1_100_n2_100_its_9999_Dist1_Asymm_bimodalvm_Dist2_vm_m1=0-0_m2_0-90_k1=2-2_k2_2-2.csv")
Figure12PD=read.csv("Results/Power_n1_40_n2_60_its_9999_Dist1_Asymm_bimodalvm_Dist2_vm_m1=0-0_m2_0-90_k1=2-2_k2_2-2.csv")
Figure12PE=read.csv( "Results/Power_n1_20_n2_100_its_9999_Dist1_Asymm_bimodalvm_Dist2_vm_m1=0-0_m2_0-90_k1=2-2_k2_2-2.csv")

list_fig12 <- list(Figure12PA, Figure12PB,Figure12PC,Figure12PD,Figure12PE)

Figure13PA=read.csv("Results/Power_n1_30_n2_30_its_9999_Dist1_asymm_trimodalvm_Dist2_vm_m1=0-0_m2_0-0_k1=0-0_k2_0-8.csv")
Figure13PB=read.csv("Results/Power_n1_60_n2_60_its_9999_Dist1_asymm_trimodalvm_Dist2_vm_m1=0-0_m2_0-0_k1=0-0_k2_0-8.csv")
Figure13PC=read.csv("Results/Power_n1_150_n2_150_its_9999_Dist1_asymm_trimodalvm_Dist2_vm_m1=0-0_m2_0-0_k1=0-0_k2_0-8.csv")
Figure13PD=read.csv("Results/Power_n1_60_n2_90_its_9999_Dist1_asymm_trimodalvm_Dist2_vm_m1=0-0_m2_0-0_k1=0-0_k2_0-8.csv")
Figure13PE=read.csv( "Results/Power_n1_30_n2_150_its_9999_Dist1_asymm_trimodalvm_Dist2_vm_m1=0-0_m2_0-0_k1=0-0_k2_0-8.csv")

list_fig13 <- list(Figure13PA, Figure13PB,Figure13PC,Figure13PD,Figure13PE)

Figure14PA=read.csv("Results/Power_n1_30_n2_30_its_9999_Dist1_asymm_trimodalvm_Dist2_vm_m1=0-0_m2_0-60_k1=2-2_k2_2-2.csv")
Figure14PB=read.csv("Results/Power_n1_60_n2_60_its_9999_Dist1_asymm_trimodalvm_Dist2_vm_m1=0-0_m2_0-60_k1=2-2_k2_2-2.csv")
Figure14PC=read.csv("Results/Power_n1_150_n2_150_its_9999_Dist1_asymm_trimodalvm_Dist2_vm_m1=0-0_m2_0-60_k1=2-2_k2_2-2.csv")
Figure14PD=read.csv("Results/Power_n1_60_n2_90_its_9999_Dist1_asymm_trimodalvm_Dist2_vm_m1=0-0_m2_0-60_k1=2-2_k2_2-2.csv")
Figure14PE=read.csv( "Results/Power_n1_30_n2_150_its_9999_Dist1_asymm_trimodalvm_Dist2_vm_m1=0-0_m2_0-60_k1=2-2_k2_2-2.csv")

list_fig14 <- list(Figure14PA, Figure14PB,Figure14PC,Figure14PD,Figure14PE)

list_all<-list(list_fig1,list_fig2,list_fig3,list_fig4,
               list_fig5,list_fig6,list_fig7,list_fig8,
               list_fig9,list_fig10,list_fig11,list_fig12,
               list_fig13,list_fig14)

nv_unimodal<-c("n = 10/10","n = 20/20","n = 50/50","n = 20/30","n = 10/50")
nv_unibi_mix<-c("n = 10/20","n = 20/40","n = 50/100","n = 20/60","n = 10/100")
nv_bimodal<-c("n = 20/20","n = 40/40","n = 100/100","n = 40/60","n = 20/100")
nv_trimodal<-c("n = 30/30","n = 60/60","n = 150/150","n = 60/90","n = 30/150")
list_nv<-list(nv_unimodal,nv_unimodal,nv_unibi_mix,nv_unimodal,
              nv_unimodal,nv_unimodal,nv_bimodal,nv_bimodal,
              nv_trimodal,nv_trimodal,nv_bimodal,nv_bimodal,
              nv_trimodal,nv_trimodal)

specs_Fig1 <- c("Figure 2.pdf","Power (VM - unimodal)","Kappa difference",1)
specs_Fig2 <- c("Figure 3.pdf","Power (VM - unimodal)","Angular difference",1)
specs_Fig3 <- c("Figure 4.pdf","Power (VM - uni/bimodal)","Kappa (for both)",1)
specs_Fig4 <- c("Figure S16.pdf","Power (VM/WSN - unimodal)","Kappa (for VM)",1)
specs_Fig5 <- c("Figure S6.pdf","Power (WSN - unimodal)","Dispersion difference",1)
specs_Fig6 <- c("Figure S11.pdf","Power (WSN - unimodal)","Angular difference",1)
specs_Fig7 <- c("Figure S7.pdf","Power (VM - axial)","Kappa difference",1)
specs_Fig8 <- c("Figure S12.pdf","Power (VM - axial)","Angular difference",1)
specs_Fig9 <- c("Figure S8.pdf","Power (VM - sym. trimodal)","Kappa difference)",1)
specs_Fig10 <- c("Figure S13.pdf","Power (VM - sym. trimodal)","Angular difference",1)
specs_Fig11 <- c("Figure S9.pdf","Power (VM - asym. bimodal)","Kappa difference",1)
specs_Fig12 <- c("Figure S14.pdf","Power (VM - asym. bimodal)","Angular difference",1)
specs_Fig13 <- c("Figure S10.pdf","Power (VM - asym. trimodal)","Kappa difference",1)
specs_Fig14 <- c("Figure S15.pdf","Power (VM - asym. trimodal)","Angular difference",1)

specs <- rbind(specs_Fig1,specs_Fig2,specs_Fig3,specs_Fig4,specs_Fig5,specs_Fig6,
               specs_Fig7,specs_Fig8,specs_Fig9,specs_Fig10,specs_Fig11,specs_Fig12,
               specs_Fig13,specs_Fig14)

#Make color scheme for the different tests 
#######color ###########
library(RColorBrewer)
qual_col_pals = brewer.pal.info[brewer.pal.info$category == 'qual',]
Col.vector= unlist(mapply(brewer.pal, qual_col_pals$maxcolors, rownames(qual_col_pals)))

#seq_brew<-seq(1,74,4)[1:18]; 
Col <- Col.vector[c(1:3,5:19)]

name_vector<- c("WatsonTwoSamp","Kuiper","MardiaWatsonWheeler","Manova","embed.circaovRfast2",
                "het.circaovRfast2","lr.circaovRfast2","WatsonLargeSample","Watson.Williams","Watson.Wheeler",
                "Raopolar","Fishers","P.Test.Assym","Wallraff","ConcTestDirectional","tang.conc","Raodispersion",
                "levenes")
name_vector_col <- paste(name_vector, "col", sep="_")

colors_plot<-mapply(function(x,y) {assign(x, y)},name_vector_col,Col)


#########Arial ##########
# library(extrafont) # in order to use Arial for figures 
# font_import() # run this line and type "yes", if fonts are not yet imported 
# loadfonts(device="pdf") 
########

for (i in 1:nrow(specs)) {
  
name_fig<-specs[,1][i]

ylab<-specs[,2][i]; xlab<-specs[,3][i]; ymax=as.numeric(specs[,4][i])

FigurexA <-list_all[[i]][[1]]; FigurexB <-list_all[[i]][[2]];FigurexC <-list_all[[i]][[3]]
FigurexD <-list_all[[i]][[4]]; FigurexE <-list_all[[i]][[5]]

x<-FigurexA$x;xmax<-max(x);xmin<-min(x);xrealmin<-min(x)
textH<-ymax+ymax*0.15
textn<-ymax+ymax*0.04


pdf(name_fig, width=8, height=2)

par(cex.axis = 0.8,cex.lab=1,mar=c(5,3,3,2),
    mfrow = c(1, 5),mgp=c(1,0.3,0),tck=-0.025,cex=0.45)
seglen=3;yesp=1;adjLet=-0.1;lineLet=1

#########prepare legends 
################Legend_1 #########

col_legen_1=colors_plot[c("WatsonTwoSamp_col","MardiaWatsonWheeler_col",
              "Manova_col","embed.circaovRfast2_col","Watson.Wheeler_col","Raopolar_col",
              "ConcTestDirectional_col",
              "levenes_col"
              )]


legend_1= function(){legend("bottom", legend=c("Watson's U� test","Large-sample Mardia-Watson-Wheeler test","MANOVA approach","Embedding approach ANOVA","Watson-Wheeler test","Rao polar test",
                                               "Concentration test","Levene's test"),
col=col_legen_1,cex = 1, lty=c(1,3,4,5,3,4,2,5),pch = c(1,1,1,1,1,2,23,23),bty="n",ncol=4,
lwd = 0.6,y.intersp = yesp,seg.len = seglen,xjust = 0, x.intersp=0.5 )}

######################## Plotting #################################
# par(oma = c(4, 1, 1, 1))

############# identical Fig 1A##### 
#First test for identical
plot(FigurexA$x,FigurexA$WatsonTwoSamp,col=colors_plot["WatsonTwoSamp_col"],type = 'b',lty=1,pch = 1,bty="n" ,
      ylab = ylab,xlab=xlab,xlim = if (xmax>5) {c(xrealmin,xmax)} else {c(xrealmin,xmax)},ylim = c(0,ymax),lwd = 0.6,xaxp = c(xrealmin, xmax,4))
lines(FigurexA$x,FigurexA$MardiaWatsonWheeler,col=colors_plot["MardiaWatsonWheeler_col"],lty=3,type = "b",pch = 1,lwd = 0.6,)
lines(FigurexA$x,FigurexA$Manova,lty=4,col=colors_plot["Manova_col"],type = "b",pch = 1,lwd = 0.6)
lines(FigurexA$x,FigurexA$embed.circaovRfast2,lty=5,col=colors_plot["embed.circaovRfast2_col"],type = "b",pch = 1,lwd = 0.6)
lines(FigurexA$x,FigurexA$Raopolar,lty=4,col=colors_plot["Raopolar_col"],type = "b",pch = 2,lwd = 0.6)
lines(FigurexA$x,FigurexA$Watson.Wheeler,col=colors_plot["Watson.Wheeler_col"],lty=3,type = "b",pch = 1,lwd = 0.6,)
lines(FigurexA$x,FigurexA$ConcTestDirectional,lty=2,col=colors_plot["ConcTestDirectional_col"],type = "b",pch = 23,lwd = 0.6)
lines(FigurexA$x,FigurexA$levenes,lty=5,col=colors_plot["levenes_col"],type = "b",pch = 23,lwd = 0.6)

#text(x=xrealmin, y=textH,labels="H0: Distributions are identical",pos=4,xpd=NA)
text(x=xrealmin, y=textn,labels=list_nv[[i]][1],pos=4,xpd=NA)
abline(h=0.05,lty=2,col="grey") 

mtext('A',side=3,cex=0.8,las=1,line=lineLet,adj=adjLet)


################ identical Fig 1B #####
#First test for identical
plot(FigurexB$x,FigurexB$WatsonTwoSamp,col=colors_plot["WatsonTwoSamp_col"],type = 'b',lty=1,pch = 1,bty="n" ,
     ylab = ylab,xlab=xlab,xlim = if (xmax>5) {c(xrealmin,xmax)} else {c(xrealmin,xmax)},ylim = c(0,ymax),lwd = 0.6,xaxp = c(xrealmin, xmax,4))
lines(FigurexB$x,FigurexB$MardiaWatsonWheeler,col=colors_plot["MardiaWatsonWheeler_col"],lty=3,type = "b",pch = 1,lwd = 0.6,)
lines(FigurexB$x,FigurexB$Manova,lty=4,col=colors_plot["Manova_col"],type = "b",pch = 1,lwd = 0.6)
lines(FigurexB$x,FigurexB$embed.circaovRfast2,lty=5,col=colors_plot["embed.circaovRfast2_col"],type = "b",pch = 1,lwd = 0.6)
lines(FigurexB$x,FigurexB$Raopolar,col=colors_plot["Raopolar_col"],lty=4,type = "b",pch = 2,lwd = 0.6,)
lines(FigurexB$x,FigurexB$Watson.Wheeler,col=colors_plot["Watson.Wheeler_col"],lty=3,type = "b",pch = 1,lwd = 0.6,)
lines(FigurexB$x,FigurexB$ConcTestDirectional,lty=2,col=colors_plot["ConcTestDirectional_col"],type = "b",pch = 23,lwd = 0.6)
lines(FigurexB$x,FigurexB$levenes,lty=5,col=colors_plot["levenes_col"],type = "b",pch = 23,lwd = 0.6)

#text(x=xrealmin, y=textH,labels="H0: Distributions are identical",pos=4,xpd=NA)
text(x=xrealmin, y=textn,labels=list_nv[[i]][2],pos=4,xpd=NA)
abline(h=0.05,lty=2,col="grey") 


mtext('B',side=3,cex=0.8,las=1,line=lineLet,adj=adjLet)

################ identical Fig 1C #####
#First test for identical
plot(FigurexC$x,FigurexC$WatsonTwoSamp,col=colors_plot["WatsonTwoSamp_col"],type = 'b',lty=1,pch = 1,bty="n" ,
     ylab = ylab,xlab=xlab,xlim = if (xmax>5) {c(xrealmin,xmax)} else {c(xrealmin,xmax)},ylim = c(0,ymax),lwd = 0.6,xaxp = c(xrealmin, xmax,4))
lines(FigurexC$x,FigurexC$MardiaWatsonWheeler,col=colors_plot["MardiaWatsonWheeler_col"],lty=3,type = "b",pch = 1,lwd = 0.6,)
lines(FigurexC$x,FigurexC$Manova,lty=4,col=colors_plot["Manova_col"],type = "b",pch = 1,lwd = 0.6)
lines(FigurexC$x,FigurexC$embed.circaovRfast2,lty=5,col=colors_plot["embed.circaovRfast2_col"],type = "b",pch = 1,lwd = 0.6)
lines(FigurexC$x,FigurexC$Raopolar,col=colors_plot["Raopolar_col"],lty=4,type = "b",pch = 2,lwd = 0.6,)
lines(FigurexC$x,FigurexC$Watson.Wheeler,col=colors_plot["Watson.Wheeler_col"],lty=3,type = "b",pch = 1,lwd = 0.6,)
lines(FigurexC$x,FigurexC$ConcTestDirectional,lty=2,col=colors_plot["ConcTestDirectional_col"],type = "b",pch = 23,lwd = 0.6)
lines(FigurexC$x,FigurexC$levenes,lty=5,col=colors_plot["levenes_col"],type = "b",pch = 23,lwd = 0.6)

#text(x=xrealmin, y=textH,labels="H0: Distributions are identical",pos=4,xpd=NA)
text(x=xrealmin, y=textn,labels=list_nv[[i]][3],pos=4,xpd=NA)
abline(h=0.05,lty=2,col="grey") 

mtext('C',side=3,cex=0.8,las=1,line=lineLet,adj=adjLet)


################ identical Fig 1D #####
#First test for identical
plot(FigurexD$x,FigurexD$WatsonTwoSamp,col=colors_plot["WatsonTwoSamp_col"],type = 'b',lty=1,pch = 1,bty="n" ,
      ylab = ylab,xlab=xlab,xlim = if (xmax>5) {c(xrealmin,xmax)} else {c(xrealmin,xmax)},ylim = c(0,ymax),lwd = 0.6,xaxp = c(xrealmin, xmax,4))
lines(FigurexD$x,FigurexD$MardiaWatsonWheeler,col=colors_plot["MardiaWatsonWheeler_col"],lty=3,type = "b",pch = 1,lwd = 0.6,)
lines(FigurexD$x,FigurexD$Manova,lty=4,col=colors_plot["Manova_col"],type = "b",pch = 1,lwd = 0.6)
lines(FigurexD$x,FigurexD$embed.circaovRfast2,lty=5,col=colors_plot["embed.circaovRfast2_col"],type = "b",pch = 1,lwd = 0.6)
lines(FigurexD$x,FigurexD$Raopolar,col=colors_plot["Raopolar_col"],lty=4,type = "b",pch = 2,lwd = 0.6,)
lines(FigurexD$x,FigurexD$Watson.Wheeler,col=colors_plot["Watson.Wheeler_col"],lty=3,type = "b",pch = 1,lwd = 0.6,)
lines(FigurexD$x,FigurexD$ConcTestDirectional,lty=2,col=colors_plot["ConcTestDirectional_col"],type = "b",pch = 23,lwd = 0.6)
lines(FigurexD$x,FigurexD$levenes,lty=5,col=colors_plot["levenes_col"],type = "b",pch = 23,lwd = 0.6)


#text(x=xrealmin, y=textH,labels="H0: Distributions are identical",pos=4,xpd=NA)
text(x=xrealmin, y=textn,labels=list_nv[[i]][4],pos=4,xpd=NA)
abline(h=0.05,lty=2,col="grey") 

mtext('D',side=3,cex=0.8,las=1,line=lineLet,adj=adjLet)

################ identical Fig 1E #####
#First test for identical
plot(FigurexE$x,FigurexE$WatsonTwoSamp,col=colors_plot["WatsonTwoSamp_col"],type = 'b',lty=1,pch = 1,bty="n" ,
     ylab = ylab,xlab=xlab,xlim = if (xmax>5) {c(xrealmin,xmax)} else {c(xrealmin,xmax)},ylim = c(0,ymax),lwd = 0.6,xaxp = c(xrealmin, xmax,4))
lines(FigurexE$x,FigurexE$MardiaWatsonWheeler,col=colors_plot["MardiaWatsonWheeler_col"],lty=3,type = "b",pch = 1,lwd = 0.6,)
lines(FigurexE$x,FigurexE$Manova,lty=4,col=colors_plot["Manova_col"],type = "b",pch = 1,lwd = 0.6)
lines(FigurexE$x,FigurexE$embed.circaovRfast2,lty=5,col=colors_plot["embed.circaovRfast2_col"],type = "b",pch = 1,lwd = 0.6)
lines(FigurexE$x,FigurexE$Raopolar,col=colors_plot["Raopolar_col"],lty=4,type = "b",pch = 2,lwd = 0.6,)
lines(FigurexE$x,FigurexE$Watson.Wheeler,col=colors_plot["Watson.Wheeler_col"],lty=3,type = "b",pch = 1,lwd = 0.6,)
lines(FigurexE$x,FigurexE$ConcTestDirectional,lty=2,col=colors_plot["ConcTestDirectional_col"],type = "b",pch = 23,lwd = 0.6)
lines(FigurexE$x,FigurexE$levenes,lty=5,col=colors_plot["levenes_col"],type = "b",pch = 23,lwd = 0.6)

#text(x=xrealmin, y=textH,labels="H0: Distributions are identical",pos=4,xpd=NA)
text(x=xrealmin, y=textn,labels=list_nv[[i]][5],pos=4,xpd=NA)
abline(h=0.05,lty=2,col="grey") 

mtext('E',side=3,cex=0.8,las=1,line=lineLet,adj=adjLet)
############################# same mean Fig 1E #####

#########Plotting the legend outside of plots#############################
par(fig = c(0, 1, 0, 1), oma = c(0, 0, 0, 0), mar = c(0, 0, 0, 0), new = TRUE)
plot(0, 0, type = "n", bty = "n", xaxt = "n", yaxt = "n")
legend_1()

dev.off()

}
```
