## Supplementary material, code and results for "Comparing two circular distributions: advice for effective implementation of statistical procedures in biology": FigureAll_Type1_plotting_Sept2020_Pascaledit_WSNreverse_LLPM.R

```
#Plotting Comp Dist

#Figure 1 (VM - identical distributions): A: (10,10), B: (20,20), C: (50,50),D: (20,30), E: (10,50), 

#load simulation data and prepare data to make it suitable for a loop
#each simulation produces 5 csv files, the basis of panels A-E
Figure1A=read.csv("Results/Type_1_error_n1_10_n2_10_its_9999_Dist1_vm_Dist2_vm_m1=0-0_m2_0-0_k1=0-8_k2_0-8.csv")
Figure1B=read.csv("Results/Type_1_error_n1_20_n2_20_its_9999_Dist1_vm_Dist2_vm_m1=0-0_m2_0-0_k1=0-8_k2_0-8.csv")
Figure1C=read.csv("Results/Type_1_error_n1_50_n2_50_its_9999_Dist1_vm_Dist2_vm_m1=0-0_m2_0-0_k1=0-8_k2_0-8.csv")
Figure1D=read.csv("Results/Type_1_error_n1_20_n2_30_its_9999_Dist1_vm_Dist2_vm_m1=0-0_m2_0-0_k1=0-8_k2_0-8.csv")
Figure1E=read.csv("Results/Type_1_error_n1_10_n2_50_its_9999_Dist1_vm_Dist2_vm_m1=0-0_m2_0-0_k1=0-8_k2_0-8.csv")

#the panels are now arranged in a list (underneath thats done for each figure)
list_fig1 <- list(Figure1A, Figure1B,Figure1C,Figure1D,Figure1E)

Figure2A=read.csv("Results/Type_1_error_n1_20_n2_20_its_9999_Dist1_Axialvm_Dist2_vm_m1=0-0_m2_0-0_k1=0-8_k2_0-8.csv")
Figure2B=read.csv("Results/Type_1_error_n1_40_n2_40_its_9999_Dist1_Axialvm_Dist2_vm_m1=0-0_m2_0-0_k1=0-8_k2_0-8.csv")
Figure2C=read.csv("Results/Type_1_error_n1_100_n2_100_its_9999_Dist1_Axialvm_Dist2_vm_m1=0-0_m2_0-0_k1=0-8_k2_0-8.csv")
Figure2D=read.csv("Results/Type_1_error_n1_40_n2_60_its_9999_Dist1_Axialvm_Dist2_vm_m1=0-0_m2_0-0_k1=0-8_k2_0-8.csv")
Figure2E=read.csv("Results/Type_1_error_n1_20_n2_100_its_9999_Dist1_Axialvm_Dist2_vm_m1=0-0_m2_0-0_k1=0-8_k2_0-8.csv")

list_fig2 <- list(Figure2A, Figure2B,Figure2C,Figure2D,Figure2E)

Figure3A=read.csv("Results/Type_1_error_n1_20_n2_20_its_9999_Dist1_Asymm_bimodalvm_Dist2_vm_m1=0-0_m2_0-0_k1=0-8_k2_0-8.csv")
Figure3B=read.csv("Results/Type_1_error_n1_40_n2_40_its_9999_Dist1_Asymm_bimodalvm_Dist2_vm_m1=0-0_m2_0-0_k1=0-8_k2_0-8.csv")
Figure3C=read.csv("Results/Type_1_error_n1_100_n2_100_its_9999_Dist1_Asymm_bimodalvm_Dist2_vm_m1=0-0_m2_0-0_k1=0-8_k2_0-8.csv")
Figure3D=read.csv("Results/Type_1_error_n1_40_n2_60_its_9999_Dist1_Asymm_bimodalvm_Dist2_vm_m1=0-0_m2_0-0_k1=0-8_k2_0-8.csv")
Figure3E=read.csv("Results/Type_1_error_n1_20_n2_100_its_9999_Dist1_Asymm_bimodalvm_Dist2_vm_m1=0-0_m2_0-0_k1=0-8_k2_0-8.csv")

list_fig3 <- list(Figure3A, Figure3B,Figure3C,Figure3D,Figure3E)


#each simulation produces 5 csv files, the basis of panels A-E
Figure4A=read.csv("Results/Type_1_error_n1_10_n2_10_its_9999_Dist1_wsn_Dist2_wsn_m1=0-0_m2_0-0_k1=1-4_k2_1-4.csv")
Figure4B=read.csv("Results/Type_1_error_n1_20_n2_20_its_9999_Dist1_wsn_Dist2_wsn_m1=0-0_m2_0-0_k1=1-4_k2_1-4.csv")
Figure4C=read.csv("Results/Type_1_error_n1_50_n2_50_its_9999_Dist1_wsn_Dist2_wsn_m1=0-0_m2_0-0_k1=1-4_k2_1-4.csv")
Figure4D=read.csv("Results/Type_1_error_n1_20_n2_30_its_9999_Dist1_wsn_Dist2_wsn_m1=0-0_m2_0-0_k1=1-4_k2_1-4.csv")
Figure4E=read.csv("Results/Type_1_error_n1_10_n2_50_its_9999_Dist1_wsn_Dist2_wsn_m1=0-0_m2_0-0_k1=1-4_k2_1-4.csv")

#the panels are now arranged in a list (underneath thats done for each figure)
list_fig4 <- list(Figure4A, Figure4B,Figure4C,Figure4D,Figure4E)

#each simulation produces 5 csv files, the basis of panels A-E
Figure5A=read.csv("Results/Type_1_error_n1_30_n2_30_its_9999_Dist1_Symm_trimodalvm_Dist2_vm_m1=0-0_m2_0-0_k1=0-8_k2_0-8.csv")
Figure5B=read.csv("Results/Type_1_error_n1_60_n2_60_its_9999_Dist1_Symm_trimodalvm_Dist2_vm_m1=0-0_m2_0-0_k1=0-8_k2_0-8.csv")
Figure5C=read.csv("Results/Type_1_error_n1_150_n2_150_its_9999_Dist1_Symm_trimodalvm_Dist2_vm_m1=0-0_m2_0-0_k1=0-8_k2_0-8.csv")
Figure5D=read.csv("Results/Type_1_error_n1_60_n2_90_its_9999_Dist1_Symm_trimodalvm_Dist2_vm_m1=0-0_m2_0-0_k1=0-8_k2_0-8.csv")
Figure5E=read.csv("Results/Type_1_error_n1_30_n2_150_its_9999_Dist1_Symm_trimodalvm_Dist2_vm_m1=0-0_m2_0-0_k1=0-8_k2_0-8.csv")

#the panels are now arranged in a list (underneath thats done for each figure)
list_fig5 <- list(Figure5A, Figure5B,Figure5C,Figure5D,Figure5E)

#each simulation produces 5 csv files, the basis of panels A-E
Figure6A=read.csv("Results/Type_1_error_n1_30_n2_30_its_9999_Dist1_asymm_trimodalvm_Dist2_vm_m1=0-0_m2_0-0_k1=0-8_k2_0-8.csv")
Figure6B=read.csv("Results/Type_1_error_n1_60_n2_60_its_9999_Dist1_asymm_trimodalvm_Dist2_vm_m1=0-0_m2_0-0_k1=0-8_k2_0-8.csv")
Figure6C=read.csv("Results/Type_1_error_n1_150_n2_150_its_9999_Dist1_asymm_trimodalvm_Dist2_vm_m1=0-0_m2_0-0_k1=0-8_k2_0-8.csv")
Figure6D=read.csv("Results/Type_1_error_n1_60_n2_90_its_9999_Dist1_asymm_trimodalvm_Dist2_vm_m1=0-0_m2_0-0_k1=0-8_k2_0-8.csv")
Figure6E=read.csv("Results/Type_1_error_n1_30_n2_150_its_9999_Dist1_asymm_trimodalvm_Dist2_vm_m1=0-0_m2_0-0_k1=0-8_k2_0-8.csv")

#the panels are now arranged in a list (underneath thats done for each figure)
list_fig6 <- list(Figure6A, Figure6B,Figure6C,Figure6D,Figure6E)

#now all the list are again arranged in a list (a list of a list)
list_all<-list(list_fig1,list_fig2,list_fig3,list_fig4,list_fig5,list_fig6)

#prepare a list for sample size labeling for each panel
nv_unimodal<-c("n = 10/10","n = 20/20","n = 50/50","n = 20/30","n = 10/50")
nv_bimodal<-c("n = 20/20","n = 40/40","n = 100/100","n = 40/60","n = 20/100")
nv_trimodal<-c("n = 30/30","n = 60/60","n = 150/150","n = 60/90","n = 30/150")
list_nv<-list(nv_unimodal,nv_bimodal,nv_bimodal,nv_unimodal,nv_trimodal,nv_trimodal)

#more text and parameters for each figure, the file name of the figure, the x-axis label, ...
# ... the y-axis label and the extent of the y-axis
specs_Fig1 <- c("Figure 1.pdf","Type 1 error (VM - unimodal)","Kappa (for both distributions)",0.6)
specs_Fig2 <- c("Figure S2.pdf","Type 1 error (VM - axial)","Kappa (for both distributions)",1)
specs_Fig3 <- c("Figure S3.pdf","Type 1 error (VM - asym. bimodal)","Kappa (for both distributions)",1)
specs_Fig4 <- c("Figure S1.pdf","Type 1 error (WSN - unimodal)","dispersion parameter (for both distributions)",1)
specs_Fig5 <- c("Figure S4.pdf","Type 1 error (VM - sym. trimodal)","Kappa (for both distributions)",1)
specs_Fig6 <- c("Figure S5.pdf","Type 1 error (VM - asym. trimodal)","Kappa (for both distributions)",1)

specs <- rbind(specs_Fig1,specs_Fig2,specs_Fig3,specs_Fig4,specs_Fig5,specs_Fig6) #binding the specs for the Figs

#Make color scheme for the different tests 
library(RColorBrewer) #load library for palette
qual_col_pals = brewer.pal.info[brewer.pal.info$category == 'qual',] 
Col.vector= unlist(mapply(brewer.pal, qual_col_pals$maxcolors, rownames(qual_col_pals))) #using palette which should give distinct colors
Col <- Col.vector[c(1:3,5:19)] #I dont use the 4th color, its a light yellow # the length of Col is equal the number of tests

#all the test names as they appear in the csv
name_vector<- c("WatsonTwoSamp","Kuiper","MardiaWatsonWheeler","Manova","embed.circaovRfast2",
                "het.circaovRfast2","lr.circaovRfast2","WatsonLargeSample","Watson.Williams","Watson.Wheeler",
                "Raopolar","Fishers","P.Test.Assym","Wallraff","ConcTestDirectional","tang.conc","Raodispersion",
                "levenes")
#adding a "col" 
name_vector_col <- paste(name_vector, "col", sep="_")

#the name plu col is now associated with a color
colors_plot<-mapply(function(x,y) {assign(x, y)},name_vector_col,Col)

# 
# #the font type 
# library(extrafont) # in order to use Arial for figures 
# font_import() # run this line and type "yes", if fonts are not yet imported 
# loadfonts(device="pdf") 


######## now the loop can start - length of the loop depends on the number of figures we want to create

for (i in 1:nrow(specs)) {
  
name_fig<-specs[,1][i]

ylab<-specs[,2][i]; xlab<-specs[,3][i]; ymax=as.numeric(specs[,4][i])

FigurexA <-list_all[[i]][[1]]; FigurexB <-list_all[[i]][[2]];FigurexC <-list_all[[i]][[3]]
FigurexD <-list_all[[i]][[4]]; FigurexE <-list_all[[i]][[5]]

x<-FigurexA$x;xmax<-max(x);xmin<-max(x)-(0.2*max(x));xrealmin<-min(x)
textH<-ymax+ymax*0.15
textn<-ymax+ymax*0.06


pdf(name_fig, width=8, height=8)

par(cex.axis = 0.8,cex.lab=1,mar=c(3,3,3,0),
    mfrow = c(5, 3),mgp=c(1,0.3,0),tck=-0.025,cex=0.45)
seglen=3;yesp=1;adjLet=-0.1;lineLet=1

#########prepare legends. I prepared them in functions which are called then inside the loops
# this approach makes the loop easier readible I think
################Legend_1 #########

col_legen_1=colors_plot[c("WatsonTwoSamp_col","Kuiper_col","MardiaWatsonWheeler_col",
              "Manova_col","embed.circaovRfast2_col",
              "het.circaovRfast2_col","lr.circaovRfast2_col","Watson.Wheeler_col")]

legend_1= function(){legend("topright", legend=c("Watson's U� test","Kuiper 2 sample test",
                                                 "Large-sample Mardia-Watson-Wheeler test","MANOVA approach",
                                                 "Embedding approach ANOVA",
                                   "Non equal concentration parameters approach ANOVA",
                                   "Log-likelihood ratio ANOVA" ,"Watson-Wheeler test"),
col=col_legen_1,cex = 1, lty=c(1:6,1,2),pch = 1,bty="n",
lwd = 0.6,y.intersp = yesp,seg.len = seglen,xjust = 0, x.intersp=0.1 )}
###########
################### Legend_2
col_legen_2=colors_plot[c("WatsonLargeSample_col","Watson.Williams_col",
                          "Raopolar_col","Fishers_col","P.Test.Assym_col")]

legend_2<-function(){legend("topright", legend=c("Watson's large-sample nonparametric test",
                                   "Watson-Williams test","Rao polar test",
                                   "Fisher's nonparametric test","P test"),
                 col=col_legen_2,cex = 1, 
                 lty=c(1:5),pch =2,bty="n",
                 lwd = 0.6,y.intersp = yesp,seg.len = seglen,xjust = 0, x.intersp=0.1)}
###########
################### Legend_3
col_legen_3=colors_plot[c("Wallraff_col","ConcTestDirectional_col","tang.conc_col","Raodispersion_col",
                "levenes_col")]

legend_3=function(){legend("topright", legend=c("Wallraff test", "Concentration test",
                         "Fisher's method","Rao dispersion test",
                         "Levene's test"),
       col=col_legen_3,cex = 1, lty=c(1:5),pch =23,bty="n",
       lwd = 0.6,y.intersp = yesp,seg.len = seglen,xjust = 0, x.intersp=0.1 )}

######################## Plotting #################################

############# identical Fig 1A##### 
#First test for identical

plot(FigurexA$x,FigurexA$WatsonTwoSamp,col=colors_plot["WatsonTwoSamp_col"],type = 'b',lty=1,pch = 1,bty="n" ,
      ylab = ylab,xlab="",xlim = if (xmax>5) {c(xrealmin,xmax)} else {c(xmax,xrealmin)},ylim = c(0,ymax),lwd = 0.6,xaxp = c(xrealmin, xmax,4))
lines(FigurexA$x,FigurexA$Kuiper,col=colors_plot["Kuiper_col"],lty=2,type = "b",pch = 1,lwd = 0.6,)
lines(FigurexA$x,FigurexA$MardiaWatsonWheeler,col=colors_plot["MardiaWatsonWheeler_col"],lty=3,type = "b",pch = 1,lwd = 0.6)
lines(FigurexA$x,FigurexA$Manova,lty=4,col=colors_plot["Manova_col"],type = "b",pch = 1,lwd = 0.6)
lines(FigurexA$x,FigurexA$embed.circaovRfast2,lty=5,col=colors_plot["embed.circaovRfast2_col"],type = "b",pch = 1,lwd = 0.6)
lines(FigurexA$x,FigurexA$het.circaovRfast2,lty=6,col=colors_plot["het.circaovRfast2_col"],type = "b",pch = 1,lwd = 0.6)
lines(FigurexA$x,FigurexA$lr.circaovRfast2,lty=1,col=colors_plot["lr.circaovRfast2_col"],type = "b",pch = 1,lwd = 0.6)
lines(FigurexA$x,FigurexA$Watson.Wheeler,col=colors_plot["Watson.Wheeler_col"],lty=2,type = "b",pch = 1,lwd = 0.6)

text(x=if (xmax>5) xrealmin else xmax, y=textH,labels="H0: Distributions are identical",pos=4,xpd=NA)
text(x=if (xmax>5) xrealmin else xmax, y=textn,labels=list_nv[[i]][1],pos=4,xpd=NA)
abline(h=0.05,lty=2,col="grey")

mtext('A',side=3,cex=0.8,las=1,line=lineLet,adj=adjLet)

legend_1()


#############same mean Fig 1A#######
######tests for mean direction 
plot(FigurexA$x,FigurexA$WatsonLargeSample,col=colors_plot["WatsonLargeSample_col"],lty=1,type = "b",pch = 2,bty="n" ,
     xlab="",ylab="",xlim = if (xmax>5) {c(xrealmin,xmax)} else {c(xmax,xrealmin)},ylim = c(0,ymax),lwd = 0.6, xaxp = c(xrealmin, xmax,4))
lines(FigurexA$x,FigurexA$Watson.Williams,col=colors_plot["Watson.Williams_col"],lty=2,type = "b",pch = 2,lwd = 0.6)
lines(FigurexA$x,FigurexA$Raopolar,col=colors_plot["Raopolar_col"],lty=3,type = "b",pch = 2,lwd = 0.6)
lines(FigurexA$x,FigurexA$Fishers,col=colors_plot["Fishers_col"],lty=4,type = "b",pch = 2,lwd = 0.6)
lines(FigurexA$x,FigurexA$P.Test.Assym,col=colors_plot["P.Test.Assym_col"],lty=5,type = "b",pch = 2,lwd = 0.6)

text(x= if (xmax>5) xrealmin else xmax, y=textH,labels="H0: Distributions have identical mean/median",pos=4,xpd=NA)
text(x=if (xmax>5) xrealmin else xmax, y=textn,labels=list_nv[[i]][1],pos=4,xpd=NA)
abline(h=0.05,lty=2,col="grey") 


legend_2()

############# concentration Fig 1A #########
########tests for concentration differences 
plot(FigurexA$x,FigurexA$Wallraff,col=colors_plot["Wallraff_col"],lty=1,type = "b",pch = 23,bty="n" ,
     xlab="",ylab="",xlim = if (xmax>5) {c(xrealmin,xmax)} else {c(xmax,xrealmin)},ylim = c(0,ymax),lwd = 0.6, xaxp = c(xrealmin, xmax,4))
lines(FigurexA$x,FigurexA$ConcTestDirectional,col=colors_plot["ConcTestDirectional_col"],lty=2,type = "b",pch = 23,lwd = 0.6)
lines(FigurexA$x,FigurexA$tang.conc,col=colors_plot["tang.conc_col"],lty=3,type = "b",pch = 23,lwd = 0.6)
lines(FigurexA$x,FigurexA$Raodispersion,col=colors_plot["Raodispersion_col"],lty=4,type = "b",pch = 23,lwd = 0.6)

lines(FigurexA$x,FigurexA$levenes,lty=5,col=colors_plot["levenes_col"],type = "b",pch = 23,lwd = 0.6)

text(x=if (xmax>5) xrealmin else xmax, y=textH,labels="H0: Distributions have identical concentration",pos=4,xpd=NA)
text(x=if (xmax>5) xrealmin else xmax, y=textn,labels=list_nv[[i]][1],pos=4,xpd=NA)
abline(h=0.05,lty=2,col="grey") 

legend_3()

################ identical Fig 1B #####
#First test for identical
plot(FigurexB$x,FigurexB$WatsonTwoSamp,col=colors_plot["WatsonTwoSamp_col"],type = 'b',lty=1,pch = 1,bty="n" ,
     ylab = ylab,xlab="",xlim = if (xmax>5) {c(xrealmin,xmax)} else {c(xmax,xrealmin)},ylim = c(0,ymax),lwd = 0.6, xaxp = c(xrealmin, xmax,4))
lines(FigurexB$x,FigurexB$Kuiper,col=colors_plot["Kuiper_col"],lty=2,type = "b",pch = 1,lwd = 0.6,)
lines(FigurexB$x,FigurexB$MardiaWatsonWheeler,col=colors_plot["MardiaWatsonWheeler_col"],lty=3,type = "b",pch = 1,lwd = 0.6,)
lines(FigurexA$x,FigurexB$Manova,lty=4,col=colors_plot["Manova_col"],type = "b",pch = 1,lwd = 0.6)
lines(FigurexA$x,FigurexB$embed.circaovRfast2,lty=5,col=colors_plot["embed.circaovRfast2_col"],type = "b",pch = 1,lwd = 0.6)
lines(FigurexA$x,FigurexB$het.circaovRfast2,lty=6,col=colors_plot["het.circaovRfast2_col"],type = "b",pch = 1,lwd = 0.6)
lines(FigurexA$x,FigurexB$lr.circaovRfast2,lty=1,col=colors_plot["lr.circaovRfast2_col"],type = "b",pch = 1,lwd = 0.6)
lines(FigurexB$x,FigurexB$Watson.Wheeler,col=colors_plot["Watson.Wheeler_col"],lty=2,type = "b",pch = 1,lwd = 0.6,)

text(x=if (xmax>5) xrealmin else xmax, y=textH,labels="H0: Distributions are identical",pos=4,xpd=NA)
text(x=if (xmax>5) xrealmin else xmax, y=textn,labels=list_nv[[i]][2],pos=4,xpd=NA)
abline(h=0.05,lty=2,col="grey") 


mtext('B',side=3,cex=0.8,las=1,line=lineLet,adj=adjLet)
############################# same mean Fig 1B #####
######tests for mean direction 
plot(FigurexB$x,FigurexB$WatsonLargeSample,col=colors_plot["WatsonLargeSample_col"],lty=1,type = "b",pch = 2,bty="n" ,
     xlab="",ylab="",xlim = if (xmax>5) {c(xrealmin,xmax)} else {c(xmax,xrealmin)},ylim = c(0,ymax),lwd = 0.6, xaxp = c(xrealmin, xmax,4))
lines(FigurexB$x,FigurexB$Watson.Williams,col=colors_plot["Watson.Williams_col"],lty=2,type = "b",pch = 2,lwd = 0.6,)

lines(FigurexB$x,FigurexB$Raopolar,col=colors_plot["Raopolar_col"],lty=3,type = "b",pch = 2,lwd = 0.6,)
lines(FigurexB$x,FigurexB$Fishers,col=colors_plot["Fishers_col"],lty=4,type = "b",pch = 2,lwd = 0.6,)
lines(FigurexB$x,FigurexB$P.Test.Assym,col=colors_plot["P.Test.Assym_col"],lty=5,type = "b",pch = 2,lwd = 0.6,)

text(x=if (xmax>5) xrealmin else xmax, y=textH,labels="H0: Distributions have identical mean/median",pos=4,xpd=NA)
text(x=if (xmax>5) xrealmin else xmax, y=textn,labels=list_nv[[i]][2],pos=4,xpd=NA)
abline(h=0.05,lty=2,col="grey") 
########################## concentration Fig 1B###############
plot(FigurexB$x,FigurexB$Wallraff,col=colors_plot["Wallraff_col"],lty=1,type = "b",pch = 23,bty="n" ,
     xlab="",ylab="",xlim = if (xmax>5) {c(xrealmin,xmax)} else {c(xmax,xrealmin)},ylim = c(0,ymax),lwd = 0.6, xaxp = c(xrealmin, xmax,4))
lines(FigurexB$x,FigurexB$ConcTestDirectional,col=colors_plot["ConcTestDirectional_col"],lty=2,type = "b",pch = 23,lwd = 0.6)
lines(FigurexB$x,FigurexB$tang.conc,col=colors_plot["tang.conc_col"],lty=3,type = "b",pch = 23,lwd = 0.6)
lines(FigurexB$x,FigurexB$Raodispersion,col=colors_plot["Raodispersion_col"],lty=4,type = "b",pch = 23,lwd = 0.6)

lines(FigurexB$x,FigurexB$levenes,lty=5,col=colors_plot["levenes_col"],type = "b",pch = 23,lwd = 0.6)

text(x=if (xmax>5) xrealmin else xmax, y=textH,labels="H0: Distributions have identical concentration",pos=4,xpd=NA)
text(x=if (xmax>5) xrealmin else xmax, y=textn,labels=list_nv[[i]][2],pos=4,xpd=NA)
abline(h=0.05,lty=2,col="grey") 

################ identical Fig 1C #####
#First test for identical
plot(FigurexC$x,FigurexC$WatsonTwoSamp,col=colors_plot["WatsonTwoSamp_col"],type = 'b',lty=1,pch = 1,bty="n" ,
     ylab = ylab,xlab="",xlim = if (xmax>5) {c(xrealmin,xmax)} else {c(xmax,xrealmin)},ylim = c(0,ymax),lwd = 0.6, xaxp = c(xrealmin, xmax,4))
lines(FigurexC$x,FigurexC$Kuiper,col=colors_plot["Kuiper_col"],lty=2,type = "b",pch = 1,lwd = 0.6,)
lines(FigurexC$x,FigurexC$MardiaWatsonWheeler,col=colors_plot["MardiaWatsonWheeler_col"],lty=3,type = "b",pch = 1,lwd = 0.6,)
lines(FigurexC$x,FigurexC$Manova,lty=4,col=colors_plot["Manova_col"],type = "b",pch = 1,lwd = 0.6)
lines(FigurexC$x,FigurexC$embed.circaovRfast2,lty=5,col=colors_plot["embed.circaovRfast2_col"],type = "b",pch = 1,lwd = 0.6)
lines(FigurexC$x,FigurexC$het.circaovRfast2,lty=6,col=colors_plot["het.circaovRfast2_col"],type = "b",pch = 1,lwd = 0.6)
lines(FigurexC$x,FigurexC$lr.circaovRfast2,lty=1,col=colors_plot["lr.circaovRfast2_col"],type = "b",pch = 1,lwd = 0.6)
lines(FigurexC$x,FigurexC$Watson.Wheeler,col=colors_plot["Watson.Wheeler_col"],lty=2,type = "b",pch = 1,lwd = 0.6,)


text(x=if (xmax>5) xrealmin else xmax, y=textH,labels="H0: Distributions are identical",pos=4,xpd=NA)
text(x=if (xmax>5) xrealmin else xmax, y=textn,labels=list_nv[[i]][3],pos=4,xpd=NA)
abline(h=0.05,lty=2,col="grey") 

mtext('C',side=3,cex=0.8,las=1,line=lineLet,adj=adjLet)

############################# same mean Fig 1C #####
######tests for mean direction 
plot(FigurexC$x,FigurexC$WatsonLargeSample,col=colors_plot["WatsonLargeSample_col"],lty=1,type = "b",pch = 2,bty="n" ,
     xlab="",ylab="",xlim = if (xmax>5) {c(xrealmin,xmax)} else {c(xmax,xrealmin)},ylim = c(0,ymax),lwd = 0.6, xaxp = c(xrealmin, xmax,4))
lines(FigurexC$x,FigurexC$Watson.Williams,col=colors_plot["Watson.Williams_col"],lty=2,type = "b",pch = 2,lwd = 0.6,)

lines(FigurexC$x,FigurexC$Raopolar,col=colors_plot["Raopolar_col"],lty=3,type = "b",pch = 2,lwd = 0.6,)
lines(FigurexC$x,FigurexC$Fishers,col=colors_plot["Fishers_col"],lty=4,type = "b",pch = 2,lwd = 0.6,)
lines(FigurexC$x,FigurexC$P.Test.Assym,col=colors_plot["P.Test.Assym_col"],lty=5,type = "b",pch = 2,lwd = 0.6,)

text(x=if (xmax>5) xrealmin else xmax, y=textH,labels="H0: Distributions have identical mean/median",pos=4,xpd=NA)
text(x=if (xmax>5) xrealmin else xmax, y=textn,labels=list_nv[[i]][3],pos=4,xpd=NA)
abline(h=0.05,lty=2,col="grey") 
########################## concentration Fig 1C###############
plot(FigurexC$x,FigurexC$Wallraff,col=colors_plot["Wallraff_col"],lty=1,type = "b",pch = 23,bty="n" ,
     xlab="",ylab="",xlim = if (xmax>5) {c(xrealmin,xmax)} else {c(xmax,xrealmin)},ylim = c(0,ymax),lwd = 0.6, xaxp = c(xrealmin, xmax,4))
lines(FigurexC$x,FigurexC$ConcTestDirectional,col=colors_plot["ConcTestDirectional_col"],lty=2,type = "b",pch = 23,lwd = 0.6)
lines(FigurexC$x,FigurexC$tang.conc,col=colors_plot["tang.conc_col"],lty=3,type = "b",pch = 23,lwd = 0.6)
lines(FigurexC$x,FigurexC$Raodispersion,col=colors_plot["Raodispersion_col"],lty=4,type = "b",pch = 23,lwd = 0.6)

lines(FigurexC$x,FigurexC$levenes,lty=5,col=colors_plot["levenes_col"],type = "b",pch = 23,lwd = 0.6)

text(x=if (xmax>5) xrealmin else xmax, y=textH,labels="H0: Distributions have identical concentration",pos=4,xpd=NA)
text(x=if (xmax>5) xrealmin else xmax, y=textn,labels=list_nv[[i]][3],pos=4,xpd=NA)
abline(h=0.05,lty=2,col="grey") 

################ identical Fig 1D #####
#First test for identical
plot(FigurexD$x,FigurexD$WatsonTwoSamp,col=colors_plot["WatsonTwoSamp_col"],type = 'b',lty=1,pch = 1,bty="n" ,
      ylab = ylab,xlab="",xlim = if (xmax>5) {c(xrealmin,xmax)} else {c(xmax,xrealmin)},ylim = c(0,ymax),lwd = 0.6, xaxp = c(xrealmin, xmax,4))
lines(FigurexD$x,FigurexD$Kuiper,col=colors_plot["Kuiper_col"],lty=2,type = "b",pch = 1,lwd = 0.6,)
lines(FigurexD$x,FigurexD$MardiaWatsonWheeler,col=colors_plot["MardiaWatsonWheeler_col"],lty=3,type = "b",pch = 1,lwd = 0.6,)
lines(FigurexD$x,FigurexD$Manova,lty=4,col=colors_plot["Manova_col"],type = "b",pch = 1,lwd = 0.6)
lines(FigurexD$x,FigurexD$embed.circaovRfast2,lty=5,col=colors_plot["embed.circaovRfast2_col"],type = "b",pch = 1,lwd = 0.6)
lines(FigurexD$x,FigurexD$het.circaovRfast2,lty=6,col=colors_plot["het.circaovRfast2_col"],type = "b",pch = 1,lwd = 0.6)
lines(FigurexD$x,FigurexD$lr.circaovRfast2,lty=1,col=colors_plot["lr.circaovRfast2_col"],type = "b",pch = 1,lwd = 0.6)
lines(FigurexD$x,FigurexD$Watson.Wheeler,col=colors_plot["Watson.Wheeler_col"],lty=2,type = "b",pch = 1,lwd = 0.6,)


text(x=if (xmax>5) xrealmin else xmax, y=textH,labels="H0: Distributions are identical",pos=4,xpd=NA)
text(x=if (xmax>5) xrealmin else xmax, y=textn,labels=list_nv[[i]][4],pos=4,xpd=NA)
abline(h=0.05,lty=2,col="grey") 

mtext('D',side=3,cex=0.8,las=1,line=lineLet,adj=adjLet)
############################# same mean Fig 1D #####
######tests for mean direction 
plot(FigurexD$x,FigurexD$WatsonLargeSample,col=colors_plot["WatsonLargeSample_col"],lty=1,type = "b",pch = 2,bty="n" ,
     xlab="",ylab="",xlim = if (xmax>5) {c(xrealmin,xmax)} else {c(xmax,xrealmin)},ylim = c(0,ymax),lwd = 0.6, xaxp = c(xrealmin, xmax,4))
#lines(FigurexD$x,FigurexD$,col=Col[7],lty=2,type = "b",pch = 2)
lines(FigurexD$x,FigurexD$Watson.Williams,col=colors_plot["Watson.Williams_col"],lty=2,type = "b",pch = 2,lwd = 0.6,)

lines(FigurexD$x,FigurexD$Raopolar,col=colors_plot["Raopolar_col"],lty=3,type = "b",pch = 2,lwd = 0.6,)
lines(FigurexD$x,FigurexD$Fishers,col=colors_plot["Fishers_col"],lty=4,type = "b",pch = 2,lwd = 0.6,)
lines(FigurexD$x,FigurexD$P.Test.Assym,col=colors_plot["P.Test.Assym_col"],lty=5,type = "b",pch = 2,lwd = 0.6,)

text(x=if (xmax>5) xrealmin else xmax, y=textH,labels="H0: Distributions have identical mean/median",pos=4,xpd=NA)
text(x=if (xmax>5) xrealmin else xmax, y=textn,labels=list_nv[[i]][4],pos=4,xpd=NA)
abline(h=0.05,lty=2,col="grey") 

########################## concentration Fig 1D###############
plot(FigurexD$x,FigurexD$Wallraff,col=colors_plot["Wallraff_col"],lty=1,type = "b",pch = 23,bty="n" ,
     xlab="",ylab="",xlim = if (xmax>5) {c(xrealmin,xmax)} else {c(xmax,xrealmin)},ylim = c(0,ymax),lwd = 0.6, xaxp = c(xrealmin, xmax,4))
lines(FigurexD$x,FigurexD$ConcTestDirectional,col=colors_plot["ConcTestDirectional_col"],lty=2,type = "b",pch = 23,lwd = 0.6)
lines(FigurexD$x,FigurexD$tang.conc,col=colors_plot["tang.conc_col"],lty=3,type = "b",pch = 23,lwd = 0.6)
lines(FigurexD$x,FigurexD$Raodispersion,col=colors_plot["Raodispersion_col"],lty=4,type = "b",pch = 23,lwd = 0.6)
lines(FigurexD$x,FigurexD$levenes,lty=5,col=colors_plot["levenes_col"],type = "b",pch = 23,lwd = 0.6)

text(x=if (xmax>5) xrealmin else xmax, y=textH,labels="H0: Distributions have identical concentration",pos=4,xpd=NA)
text(x=if (xmax>5) xrealmin else xmax, y=textn,labels=list_nv[[i]][4],pos=4,xpd=NA)
abline(h=0.05,lty=2,col="grey") 


################ identical Fig 1E #####
#First test for identical
plot(FigurexE$x,FigurexE$WatsonTwoSamp,col=colors_plot["WatsonTwoSamp_col"],type = 'b',lty=1,pch = 1,bty="n" ,
     ylab = ylab,xlab=xlab,xlim = if (xmax>5) {c(xrealmin,xmax)} else {c(xmax,xrealmin)},ylim = c(0,ymax),lwd = 0.6, xaxp = c(xrealmin, xmax,4))
lines(FigurexE$x,FigurexE$Kuiper,col=colors_plot["Kuiper_col"],lty=2,type = "b",pch = 1,lwd = 0.6,)
lines(FigurexE$x,FigurexE$MardiaWatsonWheeler,col=colors_plot["MardiaWatsonWheeler_col"],lty=3,type = "b",pch = 1,lwd = 0.6,)
lines(FigurexE$x,FigurexE$Manova,lty=4,col=colors_plot["Manova_col"],type = "b",pch = 1,lwd = 0.6)
lines(FigurexE$x,FigurexE$embed.circaovRfast2,lty=5,col=colors_plot["embed.circaovRfast2_col"],type = "b",pch = 1,lwd = 0.6)
lines(FigurexE$x,FigurexE$het.circaovRfast2,lty=6,col=colors_plot["het.circaovRfast2_col"],type = "b",pch = 1,lwd = 0.6)
lines(FigurexE$x,FigurexE$lr.circaovRfast2,lty=1,col=colors_plot["lr.circaovRfast2_col"],type = "b",pch = 1,lwd = 0.6)
lines(FigurexE$x,FigurexE$Watson.Wheeler,col=colors_plot["Watson.Wheeler_col"],lty=2,type = "b",pch = 1,lwd = 0.6,)

text(x=if (xmax>5) xrealmin else xmax, y=textH,labels="H0: Distributions are identical",pos=4,xpd=NA)
text(x=if (xmax>5) xrealmin else xmax, y=textn,labels=list_nv[[i]][5],pos=4,xpd=NA)
abline(h=0.05,lty=2,col="grey") 

mtext('E',side=3,cex=0.8,las=1,line=lineLet,adj=adjLet)
############################# same mean Fig 1E #####
######tests for mean direction 
plot(FigurexE$x,FigurexE$WatsonLargeSample,col=colors_plot["WatsonLargeSample_col"],lty=1,type = "b",pch = 2,bty="n" ,
     ylab = "",xlab=xlab,xlim = if (xmax>5) {c(xrealmin,xmax)} else {c(xmax,xrealmin)},ylim = c(0,ymax),lwd = 0.6, xaxp = c(xrealmin, xmax,4))
lines(FigurexE$x,FigurexE$Watson.Williams,col=colors_plot["Watson.Williams_col"],lty=2,type = "b",pch = 2,lwd = 0.6,)
lines(FigurexE$x,FigurexE$Raopolar,col=colors_plot["Raopolar_col"],lty=3,type = "b",pch = 2,lwd = 0.6,)
lines(FigurexE$x,FigurexE$Fishers,col=colors_plot["Fishers_col"],lty=4,type = "b",pch = 2,lwd = 0.6,)
lines(FigurexE$x,FigurexE$P.Test.Assym,col=colors_plot["P.Test.Assym_col"],lty=5,type = "b",pch = 2,lwd = 0.6,)

text(x=if (xmax>5) xrealmin else xmax, y=textH,labels="H0: Distributions have identical mean/median",pos=4,xpd=NA)
text(x=if (xmax>5) xrealmin else xmax, y=textn,labels=list_nv[[i]][5],pos=4,xpd=NA)
abline(h=0.05,lty=2,col="grey") 

########################## concentration Fig 1E###############
plot(FigurexE$x,FigurexE$Wallraff,col=colors_plot["Wallraff_col"],lty=1,type = "b",pch = 23,bty="n" ,
     ylab = "",xlab=xlab,xlim = if (xmax>5) {c(xrealmin,xmax)} else {c(xmax,xrealmin)},ylim = c(0,ymax),lwd = 0.6, xaxp = c(xrealmin, xmax,4))
lines(FigurexE$x,FigurexE$ConcTestDirectional,col=colors_plot["ConcTestDirectional_col"],lty=2,type = "b",pch = 23,lwd = 0.6)
lines(FigurexE$x,FigurexE$tang.conc,col=colors_plot["tang.conc_col"],lty=3,type = "b",pch = 23,lwd = 0.6)
lines(FigurexE$x,FigurexE$Raodispersion,col=colors_plot["Raodispersion_col"],lty=4,type = "b",pch = 23,lwd = 0.6)
lines(FigurexE$x,FigurexE$levenes,lty=5,col=colors_plot["levenes_col"],type = "b",pch = 23,lwd = 0.6)

text(x=if (xmax>5) xrealmin else xmax, y=textH,labels="H0: Distributions have identical concentration",pos=4,xpd=NA)
text(x=if (xmax>5) xrealmin else xmax, y=textn,labels=list_nv[[i]][5],pos=4,xpd=NA)
abline(h=0.05,lty=2,col="grey") 

dev.off() #thsi wrotes the plot as a pdf 

}

#Figure 1 (VM - identical mean - differing kappa): A: (10,10), B: (20,20), C: (50,50),D: (20,30), E: (10,50), 
```
