## Supplementary material, code and results for "Comparing two circular distributions: advice for effective implementation of statistical procedures in biology": Functions_JRSB_March2021.R

```
#########################################################################################################
###############P - test function - coded by Graeme Ruxton
#########################################################################################################
Pstat <- function(s1,s2){
  R1 <- rho.circular(s1)*length(s1)
  R2 <- rho.circular(s2)*length(s2)
  Rbar <- rho.circular(c(s1,s2))
  n <- length(c(s1,s2))
  R <- n*Rbar
  Rtilda <- (R1+R2)/n
  
  top1 <- (n-2)*(R1+R2-R)
  bottom1 <- n-R1-R2
  top2 <- (n-2)*(Rtilda-Rbar)*(1-Rbar)
  bottom2 <- 2*(1-Rtilda)
  return((top1/bottom1)+(top2/bottom2))
}


P.test.assym <- function(sample1,sample2){
  Pscore <- Pstat(sample1,sample2)
  n <- length(c(sample1,sample2))
  return(pf(Pscore,1,n-2,lower.tail = FALSE))}


#########################################################################################################
###############functions from Circular statistics in R (Pewsey et al. 2013)
#########################################################################################################
#YgVal Watsons Large Sample Nonparametric tests 
YgVal <- function(cdat, ndat, g) {
  N <- length(cdat) ; ndatcsum <- cumsum(ndat) 
  delhat <- 0 ; tbar <- 0
  for (k in 1:g) {
    sample <- circular(0)
    if (k==1) {low <- 0} else
      if (k > 1) {low <- ndatcsum[k-1]}
    for (j in 1:ndat[k]) { sample[j] <- cdat[j+low] }
    tm1 <- trigonometric.moment(sample, p=1)
    tm2 <- trigonometric.moment(sample, p=2)
    Rbar1 <- tm1$rho; Rbar2 <- tm2$rho ; tbar[k] <- tm1$mu
    delhat[k] <- (1-Rbar2)/(2*Rbar1*Rbar1)
  }
  dhatmax <- max(delhat) ; dhatmin <- min(delhat)
  if (dhatmax/dhatmin <= 4) {
    CP <- 0 ; SP <- 0 ; dhat0 <- 0
    for (k in 1:g) {
      CP <- CP + ndat[k]*cos(tbar[k])
      SP <- SP + ndat[k]*sin(tbar[k])
      dhat0 <- dhat0 + ndat[k]*delhat[k] 
    }
    dhat0 <- dhat0/N
    RP <- sqrt(CP*CP+SP*SP)
    Yg <- 2*(N-RP)/dhat0
    return(Yg) } 
  else if (dhatmax/dhatmin > 4) {
    CM <- 0 ; SM <- 0 ; Yg <- 0
    for (k in 1:g) {
      CM <- CM + (ndat[k]*cos(tbar[k])/delhat[k])
      SM <- SM + (ndat[k]*sin(tbar[k])/delhat[k])
      Yg <- Yg + (ndat[k]/delhat[k]) 
    }
    RM <- sqrt(CM*CM+SM*SM)
    Yg <- 2*(Yg-RM)
    return(Yg) }
}


# ################################################################################
# Calculation of Fisher's Pg test statistic for testing for a common median direction
################################################################################
MinusPiPi <- function(sample) {
  n <- length(sample)
  for (j in 1:n) {
    if (sample[j] < -pi) {sample[j] <- sample[j] + (2*pi)} else
      if (sample[j] > pi) {sample[j] <- sample[j] - (2*pi)}
  }
  return(sample)
}

PgVal <- function(cdat, ndat, g) {
  N <- length(cdat) 
  ndatcsum <- cumsum(ndat) ; gmedian <- median.circular(cdat) 
  sumterms <- 0 ; M <- 0 
  for (k in 1:g) {
    if (k==1) {low <- 0} else
      if (k > 1) {low <- ndatcsum[k-1]}
    sample <- circular(0)
    for (j in 1:ndat[k]) { sample[j] <- cdat[j+low] }
    shiftdat <- MinusPiPi(sample-gmedian) ; m <- length(shiftdat[shiftdat<0]) ; M <- M+m
    sumterms <- sumterms + m*m/ndat[k]
  }
  term1 <- ((N*N)/(M*(N-M))); term2 <- (N*M)/(N-M) ; Pg <- term1*sumterms-term2
  return(Pg)
}
#Fisher's large-sample test for a common concentration of von Mises distributions
################################################################################
dValues <- function(cdat, ndat, g) {
  N <- length(cdat) ; ndatcsum <- cumsum(ndat) ; dval <- 0 
  for (k in 1:g) {
    sample <- circular(0) 
    if (k==1) {low <- 0} else
      if (k > 1) {low <- ndatcsum[k-1]}
    for (j in 1:ndat[k]) { sample[j] <- cdat[j+low] }
    tm1 <- trigonometric.moment(sample, p=1) ; tbar <- tm1$mu
    dvalk <- abs(sin(sample-tbar))
    dval <- c(dval, dvalk)
  }
  dval <- dval[-1]
  return(dval)
}

FgVal <- function(dvals, ndat, g) {
  N <- length(dvals) ; ndatcsum <- cumsum(ndat) 
  sum1 <- 0 ; sum2 <- 0 ; dk <- 0 ; dbar <- 0 ; gdbar <- 0 
  for (k in 1:g) {
    sample <- circular(0) 
    if (k==1) {low <- 0} else
      if (k > 1) {low <- ndatcsum[k-1]}
    for (j in 1:ndat[k]) { dk[j] <- dvals[j+low] }
    dbar[k] <- sum(dk)/ndat[k]
    sum2 <- sum2 + sum((dk-dbar[k])**2)
    gdbar <- gdbar+ndat[k]*dbar[k]
  }
  gdbar <- gdbar/N
  for (k in 1:g) { sum1 <- sum1 + ndat[k]*(dbar[k]-gdbar)**2 }
  Fg <- (N-g)*sum1/((g-1)*sum2)
  return(Fg)
}
####################
################################################################################
# Large-sample Mardia-Watson-Wheeler test for a common distribution
################################################################################
CosSinUniScores <- function(cdat) {
  N <- length(cdat)
  ranks <- rank(cdat, ties.method="random")
  CosUniScores <- cos(ranks*2*pi/N)
  SinUniScores <- sin(ranks*2*pi/N)
  return(list(CosUniScores, SinUniScores))
}

WgVal <- function(CSUScores, ndat, g) {
  CosUScores <- CSUScores[[1]] ; SinUScores <- CSUScores[[2]]
  N <- length(CosUScores) ; ndatcsum <- cumsum(ndat)
  Wg <- 0
  for (k in 1:g) {
    CosUScoresk <- 0 ; SinUScoresk <- 0
    if (k==1) {low <- 0} else
      if (k > 1) {low <- ndatcsum[k-1]}
    for (j in 1:ndat[k]) { 
      CosUScoresk[j] <- CosUScores[j+low] ; SinUScoresk[j] <- SinUScores[j+low] }
    sumCkSq <- (sum(CosUScoresk))**2 ; sumSkSq <- (sum(SinUScoresk))**2
    Wg <- Wg+(sumCkSq+sumSkSq)/ndat[k]
  }
  Wg <- 2*Wg ; return(Wg)
}


###########circular plotting wrapper (for unimodal distributions) - from Pail et al. 2020 Herpetozoa

wrapper_unimodal_plot = function (x,x_col,labels = NULL) {
  
  ori=circular(x);
  p=rayleigh.test(ori);p=p$p.value
  
  rh=rho.circular(ori);me=circular(mean.circular(ori))
  plot.circular(ori,stack = T,axes=FALSE,template = "geographics", units = "degrees",start.sep = start.sep ,
                sep=sep,shrink=shrink, cex = cexD,pch=16, tol =0,col=x_col)
  axis.circularLL(at=circular(rad(c(90))), labels,tcl.text=tcltext,
                  tick=TRUE,
                  tcl=tclength,
                  cex = cexlabel)
  arrows.circular(x=me,y=rh,lwd=lwdarrow,length=lengtharrow,zero = rad(90),code = 2,rotation ="clock")
  
  if (p<0.05) (plot_bootstrap_CI_inRad_outDeg(ori)) else (NA)
}


########### wrapper for bootstrap CI
plot_bootstrap_CI_inRad_outDeg =function (x) 
{bootci <- mle.vonmises.bootstrap.ci(x, alpha = 0.05,reps = 10000, control.circular = list())
ci2_5 <-(bootci[["mu.ci"]][["2.5%"]])
ci97_5 <-(bootci[["mu.ci"]][["97.5%"]])

arrows.circular(x=ci2_5,y=1,lwd=1,lty=2, rotation = "clock",zero = rad(90),code = 0) 
arrows.circular(x=ci97_5,y=1,lwd=1,lty=2, rotation = "clock",zero = rad(90),code = 0) 
}


#####################axis.circularLL - basically axis circular without center cross 


axis.circularLL =function (at = NULL, labels = NULL, units = NULL, template = NULL, 
                           modulo = NULL, zero = NULL, rotation = NULL, tick = TRUE, 
                           lty, lwd, cex, col, font, tcl = 0.025, tcl.text = 0.125, 
                           digits = 2) 
{
  if (missing(cex)) 
    cex <- par("cex.axis")
  if (missing(col)) 
    col <- par("col.axis")
  if (missing(font)) 
    font <- par("font.axis")
  if (missing(lty)) 
    lty <- par("lty")
  if (missing(lwd)) 
    lwd <- par("lwd")
  if (is.null(at)) {
    if (is.null(template) | template == "none" | template == 
        "geographics") {
      at <- circular(c(0, pi/2, pi, 3/2 * pi), rotation = rotation, 
                     zero = zero)
    }
    else if (template == "clock24") {
      at <- circular(seq(0, 23), units = "hours", rotation = rotation, 
                     zero = zero)
      units <- "hours"
    }
    else if (template == "clock12") {
      at <- circular(seq(0, 11), units = "hours", rotation = rotation, 
                     zero = zero)
      units <- "hours"
    }
  }
  at <- na.omit(at)
  atcircularp <- attr(as.circular(at), "circularp")
  type <- atcircularp$type
  if (is.null(modulo)) 
    modulo <- atcircularp$modulo
  if (is.null(units)) 
    units <- atcircularp$units
  if (is.null(template)) 
    template <- atcircularp$template
  if (template == "geographics" | template == "clock24") {
    zero <- pi/2
    rotation <- "clock"
  }
  else if (template == "clock12") {
    zero <- pi/2
    rotation <- "clock"
  }
  else {
    if (is.null(zero)) 
      zero <- atcircularp$zero
    if (is.null(rotation)) 
      rotation <- atcircularp$rotation
  }
  atasis <- at
  attr(atasis, "circularp") <- attr(atasis, "class") <- NULL
  attext <- atasis/pi
  if (modulo == "2pi") {
    if (units == "radians") {
      atasis <- atasis%%(2 * pi)
    }
    else if (units == "degrees") {
      atasis <- atasis%%(360)
    }
    else if (units == "hours") {
      atasis <- atasis%%(24)
    }
    attext <- attext%%2
  }
  else if (modulo == "pi") {
    if (units == "radians") {
      atasis <- atasis%%pi
    }
    else if (units == "degrees") {
      atasis <- atasis%%180
    }
    else if (units == "hours") {
      atasis <- atasis%%12
    }
    attext <- attext%%1
  }
  attext <- round(attext, digits = digits)
  if (template == "clock12") 
    at <- 2 * at
  at <- conversion.circular(at, units = "radians", modulo = "2pi", 
                            zero = 0, rotation = "counter")
  attr(at, "circularp") <- attr(at, "class") <- NULL
  if (is.null(labels)) {
    if (length(atasis) == 4 && all(atasis == c(0, pi/2, 
                                               pi, 3/2 * pi))) {
      if (template == "geographics") {
        labels <- c("N", "E", "S", "W")
      }
      else {
        if (units == "radians") {
          labels <- c("0", expression(frac(pi, 2)), 
                      expression(pi), expression(frac(3 * pi, 
                                                      2)))
        }
        else if (units == "degrees") {
          labels <- c("0", "90", "180", "270")
        }
      }
    }
    else if (length(atasis) == 12 && all(atasis == 0:11) && 
             template == "clock12") {
      labels <- c("0/12", "1", "2", "3", "4", "5", "6", 
                  "7", "8", "9", "10", "11")
    }
    else if (length(atasis) == 24 && all(atasis == 0:23) && 
             template == "clock24") {
      labels <- c("0/24", "1", "2", "3", "4", "5", "6", 
                  "7", "8", "9", "10", "11", "12", "13", "14", 
                  "15", "16", "17", "18", "19", "20", "21", "22", 
                  "23")
    }
    else if (units == "radians") {
      labels <- as.character(round(atasis, digits = digits))
    }
    else if (units == "degrees") {
      labels <- as.character(round(atasis, digits = digits))
    }
    else if (units == "hours") {
      labels <- as.character(round(atasis, digits = digits))
    }
  }
  if (!is.null(labels) && length(at) != length(labels)) 
    stop("'at' and 'labels' must have the same length")
  AxisCircularRadLL(at, units, labels, attext, tick, tcl, tcl.text, 
                    cex, col, lty, lwd)
}

AxisCircularRadLL = function (at, units, labels, attext, tick, tcl, tcl.text, cex, 
                              col, lty, lwd) 
{
  r <- 1 + tcl * c(-1/2, 1/2)
  r.l <- 1 - tcl.text
  z <- cos(at)
  y <- sin(at)
  for (i in 1:length(at)) {
    if (tick) {
      lines.default(z[i] * r, y[i] * r, col = col, lty = lty, 
                    lwd = lwd)
    }
    if (is.null(labels)) {
      labeltext <- substitute(at * pi, list(at = attext[i]))
    }
    else {
      labeltext <- labels[i]
    }
    text.default(z[i] * r.l, y[i] * r.l, labeltext, cex = cex, 
                 col = col)
  }
  
}
```
