## Supplementary material, code and results for "Comparing two circular distributions: advice for effective implementation of statistical procedures in biology": CompDist-2AsymmTrimVonMises-All_N_foreach_new.R

```
# Code to test which tests are more powerful in detecting differences between distributions 
library(kuiper.2samp)
library(lawstat)
library(circular)
library(compiler)
library(Directional)
library(NPCirc)
library(CircStats)
library(Rfast2)
library(foreach)
library(doParallel)
options("scipen" = 10)
enableJIT(3)

source("Functions_JRSB_March2021.R") # only works like this if the functions are in the working directory, otherwise change path to working directory 
options("scipen" = 10)
enableJIT(3)

nvseq=c(10,20,50,20,10)*3
nvseq2=c(10,20,50,30,50)*3
nv1=c(nvseq,nvseq,nvseq)
nv2=c(nvseq2,nvseq2,nvseq2)
n_matrix = cbind(nv1,nv2)

kseq=c(0,0.25,0.5,1,2,4,8)
lk=length(kseq)
k0=rep(0,7)
k2=rep(2,7)

mseq= c(rad(0),rad(30),rad(60),rad(90),rad(120),rad(150),rad(180))/3
lm=length(mseq)
m0=rep(0,7)

k_list_1=list(kseq,kseq,kseq,kseq,kseq,k0,k0,k0,k0,k0,k2,k2,k2,k2,k2)
k_list_2=list(kseq,kseq,kseq,kseq,kseq,kseq,kseq,kseq,kseq,kseq,k2,k2,k2,k2,k2)
m_list_1=list(m0,m0,m0,m0,m0,m0,m0,m0,m0,m0,m0,m0,m0,m0,m0)
m_list_2=list(m0,m0,m0,m0,m0,m0,m0,m0,m0,m0,mseq,mseq,mseq,mseq,mseq)
 

l=length(kseq)
type_vm=c("vm","vm","vm","vm","vm") 
type_matrix_1 = c(type_vm,type_vm,type_vm)  
type_matrix_2 = c(type_vm,type_vm,type_vm)  

Type_1_errorn=c("Type_1_error","Type_1_error","Type_1_error","Type_1_error","Type_1_error")
Powern=c("Power","Power","Power","Power","Power")

Name=c(Type_1_errorn,Powern,Powern)

########### set number of iterations, Sample size mean and kappa (or a sequence of any of that)
rans = 9999 #iterations for  distribution 


for (i in 1:length(nv1)) {
  

nv1 = n_matrix[,1][i]
nv2=n_matrix[,2][i]
m1seq=m_list_1[[i]]
m2seq=m_list_2[[i]]
k1seq=k_list_1[[i]]
k2seq=k_list_2[[i]]


typeDist1 =type_matrix_1[i]
typeDist2 = type_matrix_2[i]


# make two groups for the test where the data needs to b sturctured in the same column
Group1 <-(as.matrix( rep((as.matrix((strrep(c("A"), 1)))),nv1),dim=c(nv1, 1)))
Group2 <- (as.matrix( rep((as.matrix((strrep(c("B"), 1)))),nv2),dim=c(nv2, 1)))
GroupsAB<-as.data.frame(c(Group1[,1], Group2[,1]))

#setup parallel backend to use many processors
cores=detectCores()
cl <- parallel::makeCluster(cores[1]-1, setup_strategy = "sequential")
registerDoParallel(cl)


Power.sample<-foreach(e = 1:l,.combine=rbind,.packages=c("kuiper.2samp","lawstat","circular",
                                                 "Directional","NPCirc",
                                                 "CircStats","Rfast2")) %dopar% {
  m1 = m1seq[e]
  m2 = m2seq[e]                                       
  k1 = k1seq[e]
  k2 = k2seq[e]

  #First distribution 
  
  Distribution1 <- as.data.frame(sapply(1:rans, function(x) rcircmix(nv1,model = NULL, 
                                                                dist = c(typeDist1,typeDist1,typeDist1),
                                                                param=list(p=c(0.333,0.333,0.333), 
                                                                           mu=c(m1,m1-rad(90),m1-rad(200)), 
                                                                           con=c(k1,k1,k1)))),units = 'radians')
  
  ###### set the 2nd distribution 
  
  
  Distribution2 <- as.data.frame(sapply(1:rans, function(x) rcircmix(nv2,model = NULL, 
                                                                dist = c(typeDist2,typeDist2,typeDist2),
                                                                param=list(p=c(0.333,0.333,0.333), 
                                                                   mu=c(m2,m2-rad(90),m2-rad(200)), 
                                                     con=c(k2,k2,k2)))),units = 'radians')
  ######
  ##watson2SampleTest
 watson1 = mapply(function(x,y) {watson.two.test(x,y)},Distribution1,Distribution2)
 watson1 <- array(as.matrix(unlist(watson1)), dim=c(5, rans))
 watson1<- watson1[1,]  # critical valueCritical Value: 0.187 
 
 ##kuiper

 kuiper1 = mapply(function(x,y) {kuiper.2samp(x, y)},Distribution1,Distribution2)
 kuiper1 <- array(as.matrix(unlist(kuiper1)), dim=c(2, rans))
 kuiper1<- kuiper1[2,]
 
 ##P test Assym 
 
 P.Test.Assym_P = mapply(function(x,y) {P.test.assym(x, y)},Distribution1,Distribution2)
 

  #Raos Tests for Homogeneity
  rao1 = mapply(function(x,y) {capture.output(rao.homogeneity(list(x, y),alpha=0.05))},Distribution1,Distribution2)
  raopolarP<- rao1[9,]
  raodispersionP<- rao1[16,]

 #####preparing sample in the way I need it for the tests in the Pewsey et al. - and getting ndat and g
  DistGroups<-mapply(function(x,y) {c(x, y)},Distribution1,Distribution2)
  ndat <- c(nv1, nv2)
  g <- length(ndat)

    #Watsons Large Sample Nonparametric test
  WatsonLargeP<-(apply(DistGroups,2, FUN=function(x) pchisq(YgVal(x, ndat, g), g-1, lower.tail=F)))
  
 
  #Fisher's Pg test statistic for testing for a common median direction
  FishersP<-(apply(DistGroups,2, FUN=function(x) pchisq(PgVal(x, ndat, g), g-1, lower.tail=F)))
  

  #Fisher's large-sample test for a common concentration of von Mises distributions
  N<-nv1+nv2
  FishersLargeP<-(apply(DistGroups,2, FUN=function(x) pf(FgVal(dValues(x, ndat, g), ndat, g), g-1, N-g, lower.tail=F)))

  # Large-sample Mardia-Watson-Wheeler test for a common distribution
  MardiaWatsonWheelerP<-(apply(DistGroups,2, FUN=function(x) pchisq(WgVal(CosSinUniScores(x), ndat, g), 2*(g-1), lower.tail=F))) # WHY 2*(g-1)??? in other code  only g-1

    ###########
  #################
  #sort distributions underneath each other, is already done for Pewsey - doubel check that! 
  #  DistGroups <-mapply(function(x,y) {c(x, y)},Distribution1,Distribution2)
  #Levenes # 
  LevenesRES <- apply(DistGroups,2,FUN=function(x) lawstat::levene.test(x,GroupsAB[,1]))
  LeveneP <- sapply(c(1:rans),function(x) LevenesRES[[x]][["p.value"]])
  
  #FromDirectionalPackage  Test of equality of the concentration parameters for circular data 
  tangconcDir <- apply(DistGroups,2,FUN=function(x) tang.conc(x,as.factor(GroupsAB[,1]),rads=TRUE))
  tangconcDirP<- tangconcDir[2,] 
  
  #FromDirectionalPackage  Test of equality of the concentration parameters for circular data from Mardia, K. V. and Jupp, P. E. (2000). Directional statistics. Chicester: John Wiley & Sons.
  conctestDir <- apply(DistGroups,2,FUN=function(x) conc.test(x,as.factor(GroupsAB[,1]),rads=TRUE))
  conctestDirP<-as.character(sapply(c(1:rans),function(x) conctestDir[[x]][["res"]][["p-value"]]))
  
  #embed.circaov from Rfast2 (originally from Directional)
  embed.circaovRfast2 <- apply(DistGroups,2,FUN=function(x) Rfast2::embed.circaov(x,as.factor(GroupsAB[,1])))
  embed.circaovRfast2p<-embed.circaovRfast2[2,]
  
 
  #hRfast2::het.circaov (originally from Directional)
  het.circaovRfast2 <- apply(DistGroups,2,FUN=function(x) Rfast2::het.circaov(x,as.factor(GroupsAB[,1])))
  het.circaovRfast2p<-het.circaovRfast2[2,]
  
  #hRfast2::het.circaov (originally from Directional)
  lr.circaovRfast2 <- apply(DistGroups,2,FUN=function(x) Rfast2::lr.circaov(x,as.factor(GroupsAB[,1])))
  lr.circaovRfast2p<-lr.circaovRfast2[2,]
  
  #Watson wheeler test! Test assumes von Mises
  watson.wheelerRES <- apply(DistGroups,2,FUN=function(x) watson.wheeler.test(x,GroupsAB[,1]))
  watson.wheelerP <- sapply(c(1:rans),function(x) watson.wheelerRES[[x]][["p.value"]])
  
  #Watson william test! Test assumes von Mises and equal kappa
  watson.williamsRES <- apply(DistGroups,2,FUN=function(x) watson.williams.test(x,GroupsAB[,1]))
  watson.williamsP<- sapply(c(1:rans),function(x) watson.williamsRES[[x]][["p.value"]])
  
  #Wallraff test!
  wallraff <- apply(DistGroups,2,FUN=function(x) wallraff.test(x,GroupsAB[,1],ref=NULL))
  wallraffP<- sapply(c(1:rans),function(x) wallraff[[x]][["p.value"]])
  
  ##MANOVA approach 
   funManCirc<-function(x) summary(manova(cbind(cos(x),sin(x)) ~ GroupsAB[,1]))
   MANOVA_approach <- apply(DistGroups,2, FUN=funManCirc)
   MANOVA_P <- sapply(c(1:rans),function(x) MANOVA_approach[[x]][["stats"]][1,6])
   
   temp <- data.frame(matrix(nrow = 1, ncol = 19))
   names(temp) <- c('tang.conc','levenes','WatsonTwoSamp','Watson.Wheeler',
                            "Watson.Williams",'Wallraff', 'Raodispersion', 
                            "Raopolar","ConcTestDirectional", "WatsonLargeSample",
                           "Fishers","FishersLarge", "MardiaWatsonWheeler", 
                            "Kuiper","Manova","P.Test.Assym","embed.circaovRfast2",
                    "het.circaovRfast2","lr.circaovRfast2")
   
  #####Calculate power in temp file 
  temp$levenes<- (sum(1*(LeveneP<0.05)))/rans
  temp$tang.conc<- (sum(1*(tangconcDirP<0.05)))/rans
  temp$WatsonTwoSamp<- (sum(1*(watson1>0.178)))/rans
  temp$Watson.Wheeler<- (sum(1*(watson.wheelerP<0.05)))/rans
  temp$Watson.Williams<- (sum(1*(watson.williamsP<0.05)))/rans
  temp$Wallraff<- (sum(1*(wallraffP<0.05)))/rans
  temp$Raodispersion <- sum(1*(raodispersionP == "Reject null hypothesis of equal dispersions "))/rans
  temp$Raopolar <- sum(1*(raopolarP != "Do not reject null hypothesis of equal polar vectors "))/rans
  temp$ConcTestDirectional<-(sum(1*(conctestDirP<0.05)))/rans
  temp$embed.circaovRfast2<-(sum(1*(embed.circaovRfast2p<0.05)))/rans
 temp$het.circaovRfast2<-(sum(1*(het.circaovRfast2p<0.05)))/rans
  temp$lr.circaovRfast2<-(sum(1*(lr.circaovRfast2p<0.05)))/rans
  temp$WatsonLargeSample<-(sum(1*(WatsonLargeP<0.05)))/rans
  temp$Fishers<-(sum(1*(FishersP<0.05)))/rans
  temp$FishersLarge<-(sum(1*(FishersLargeP<0.05)))/rans
  temp$MardiaWatsonWheeler<-(sum(1*(MardiaWatsonWheelerP<0.05)))/rans
  temp$Kuiper<- (sum(1*(kuiper1<0.05)))/rans
  temp$Manova<- (sum(1*(MANOVA_P<0.05)))/rans
  temp$P.Test.Assym<- (sum(1*(P.Test.Assym_P<0.05)))/rans

    temp
} #e
stopCluster(cl)

Power.sample$x=if(max(k2seq)>min(k2seq)) k2seq else deg(m2seq)

text = paste(Name[i],"_n1_",nv1,"_n2_",nv2,"_its_",rans,"_Dist1_","asymm_trimodal",typeDist1,"_Dist2_",typeDist2,
             "_m1=",deg(min(m1seq)),"-",deg(max(m1seq)),"_m2_",deg(min(m2seq)),"-",deg(max(m2seq)),
             "_k1=",min(k1seq),"-",max(k1seq),"_k2_",min(k2seq),"-",max(k2seq),".csv",sep="")
write.csv(Power.sample, file = text,row.names=FALSE )

}#i
```
