## Supplementary material, code and results for "Comparing two circular distributions: advice for effective implementation of statistical procedures in biology": CompDist-2VonMises-All_N_foreach_new.R

```
# Code to test which tests are more powerful in detecting differences between distributions 
library(kuiper.2samp)
library(lawstat)
library(circular)
library(compiler)
library(Directional)
library(NPCirc)
library(CircStats)
library(Rfast2)
library(foreach)
library(doParallel)
options("scipen" = 10)
enableJIT(3)

source("Functions_JRSB_March2021.R") # only works like this if the functions are in the working directory, otherwise change path to working directory 
options("scipen" = 10)
enableJIT(3)

#setting sample size for dist 1 - in two steps to reduce typing effort
nvseq=c(10,20,50,20,10)
nv1=c(nvseq,nvseq,nvseq) 

#setting sample size for dist 2
nvseq2=c(10,20,50,30,50) 
nv2=c(nvseq2,nvseq2,nvseq2)

#combine sample sizes in matrix to make calling in the loop practical
n_matrix = cbind(nv1,nv2)

#preparing certain vectors of concentration parameters and direction - all 7 elements long (they need to be same length) 

kseq=c(0,0.25,0.5,1,2,4,8) #standard kappa sequence for plot 

k0=rep(0,7)
k2=rep(2,7)

mseq= c(rad(0),rad(30),rad(60),rad(90),rad(120),rad(150),rad(180))
m0=rep(0,7)

#this would be needed of lengths differ, at the present it doesn't matter which l is used 
lk=length(kseq) 
lm=length(mseq)
l=lk # this one is in this script used for the inner loop

#preparing the sequences in a list so they can be called in inner loop 
k_list_1=list(kseq,kseq,kseq,kseq,kseq,k0,k0,k0,k0,k0,k2,k2,k2,k2,k2)
k_list_2=list(kseq,kseq,kseq,kseq,kseq,kseq,kseq,kseq,kseq,kseq,k2,k2,k2,k2,k2)
m_list_1=list(m0,m0,m0,m0,m0,m0,m0,m0,m0,m0,m0,m0,m0,m0,m0)
m_list_2=list(m0,m0,m0,m0,m0,m0,m0,m0,m0,m0,mseq,mseq,mseq,mseq,mseq)
 

#defining the distribution type, there are always 5 combinations of sample sizes 
#therefore I prepared "chunks" of distribution and then combined them, decreasing typing effort 
type_vm=c("vm","vm","vm","vm","vm") 
type_matrix_1 = c(type_vm,type_vm,type_vm)  
type_matrix_2 = c(type_vm,type_vm,type_vm)  

#again in chunks of 5: naming the file that is created (see text for write.csv)
Type_1_errorn=c("Type_1_error","Type_1_error","Type_1_error","Type_1_error","Type_1_error")
Powern=c("Power","Power","Power","Power","Power")

#again combining the chunks 
Name=c(Type_1_errorn,Powern,Powern)

########### set number of iterations, Sample size mean and kappa (or a sequence of any of that)
rans = 9999 #iterations for  distribution

#starting the outer loop, it loops through the sample size vector, taking the respective vectors (of length 7) from the respective list
#the outer loop is stored one csv file, this means one panel in the plot. 
for (i in 1:length(nv1)) {

#grabing the correct vectros for the simulation from the list
nv1 = n_matrix[,1][i]
nv2=n_matrix[,2][i]
m1seq=m_list_1[[i]]
m2seq=m_list_2[[i]]
k1seq=k_list_1[[i]]
k2seq=k_list_2[[i]]


typeDist1 =type_matrix_1[i]
typeDist2 = type_matrix_2[i]


# make AB data.frame for the tests where the data needs to be structured in the same column
#The data frame consists of A and Bs arranged underneath each other indicating the groups (like a "treatment" column) 
Group1 <-(as.matrix( rep((as.matrix((strrep(c("A"), 1)))),nv1),dim=c(nv1, 1)))
Group2 <- (as.matrix( rep((as.matrix((strrep(c("B"), 1)))),nv2),dim=c(nv2, 1)))
GroupsAB<-as.data.frame(c(Group1[,1], Group2[,1]))

#setup parallel back-end to use many processors
cores=detectCores()
cl <- parallel::makeCluster(cores[1]-1, setup_strategy = "sequential") #used "sequential" otherwise it crashed for me 
registerDoParallel(cl)

#start the inner loop. Here I use a foreach instead of for. It splits the jobs on the number of cores set up. 
#Parallelization is done with "%dopar%". The packages used in the foreach loop must be passed to the loop.
#they are not grabbed automatically from the environment.
#the results of each of the loops are combined (with rbind) in the file Power.sample
Power.sample<-foreach(e = 1:l,.combine=rbind,.packages=c("kuiper.2samp","lawstat","circular",
                                                 "Directional","NPCirc",
                                                 "CircStats","Rfast2")) %dopar% {

#take correct values for mean direction as well as the concentration parameter from the vector
 m1 = m1seq[e] 
  m2 = m2seq[e]                                       
  k1 = k1seq[e]
  k2 = k2seq[e]

  #First distribution generated rans times, the "units=" could be deleted. Its just here to remind me what it is, but doesn't do anything.
  
  Distribution1 <- as.data.frame(sapply(1:rans, function(x) rcircmix(nv1,model = NULL, 
                                                                dist = typeDist1,
                                                                param=list(p=1, 
                                                                           mu=m1, 
                                                                           con=k1))))
  
  ###### set up the 2nd distribution 
  Distribution2 <- as.data.frame(sapply(1:rans, function(x) rcircmix(nv2,model = NULL, 
                                                                dist = typeDist2,
                                                                param=list(p=1, 
                                                                   mu=m2, 
                                                     con=k2))))
  
  ###### underneath we apply each of the tests used on the generated distribution 
  #using mapply (if the groups are arranged in separate column) and apply (if they are arranged underneath each other)
  
  ##watson2SampleTest
 watson1 = mapply(function(x,y) {watson.two.test(x,y)},Distribution1,Distribution2)
 watson1 <- array(as.matrix(unlist(watson1)), dim=c(5, rans))
 watson1<- watson1[1,]  # critical valueCritical Value: 0.187 
 
 ##kuiper

 kuiper1 = mapply(function(x,y) {kuiper.2samp(x, y)},Distribution1,Distribution2)
 kuiper1 <- array(as.matrix(unlist(kuiper1)), dim=c(2, rans))
 kuiper1<- kuiper1[2,]
 
 ##P test Assym 
 P.Test.Assym_P = mapply(function(x,y) {P.test.assym(x, y)},Distribution1,Distribution2)
 
  #Raos Tests for Homogeneity
  rao1 = mapply(function(x,y) {capture.output(rao.homogeneity(list(x, y),alpha=0.05))},Distribution1,Distribution2)
  raopolarP<- rao1[9,]
  raodispersionP<- rao1[16,]

 #####preparing sample in the way I need it for the tests in the Pewsey et al. - and getting ndat and g
  DistGroups<-mapply(function(x,y) {c(x, y)},Distribution1,Distribution2)
  ndat <- c(nv1, nv2)
  g <- length(ndat)

  #Watsons Large Sample Nonparametric test
  WatsonLargeP<-(apply(DistGroups,2, FUN=function(x) pchisq(YgVal(x, ndat, g), g-1, lower.tail=F)))
  
 
  #Fisher's Pg test statistic for testing for a common median direction
  FishersP<-(apply(DistGroups,2, FUN=function(x) pchisq(PgVal(x, ndat, g), g-1, lower.tail=F)))
  

  #Fisher's large-sample test for a common concentration of von Mises distributions
  N<-nv1+nv2
  FishersLargeP<-(apply(DistGroups,2, FUN=function(x) pf(FgVal(dValues(x, ndat, g), ndat, g), g-1, N-g, lower.tail=F)))

  # Large-sample Mardia-Watson-Wheeler test for a common distribution
  MardiaWatsonWheelerP<-(apply(DistGroups,2, FUN=function(x) pchisq(WgVal(CosSinUniScores(x), ndat, g), 2*(g-1), lower.tail=F))) # WHY 2*(g-1)??? in other code  only g-1

  #Levenes # 
  LevenesRES <- apply(DistGroups,2,FUN=function(x) lawstat::levene.test(x,GroupsAB[,1]))
  LeveneP <- sapply(c(1:rans),function(x) LevenesRES[[x]][["p.value"]])
  
  #FromDirectionalPackage  Test of equality of the concentration parameters for circular data 
  tangconcDir <- apply(DistGroups,2,FUN=function(x) tang.conc(x,as.factor(GroupsAB[,1]),rads=TRUE))
  tangconcDirP<- tangconcDir[2,] 
  
  #FromDirectionalPackage  Test of equality of the concentration parameters for circular data from Mardia, K. V. and Jupp, P. E. (2000). Directional statistics. Chicester: John Wiley & Sons.
  conctestDir <- apply(DistGroups,2,FUN=function(x) conc.test(x,as.factor(GroupsAB[,1]),rads=TRUE))
  conctestDirP<-as.character(sapply(c(1:rans),function(x) conctestDir[[x]][["res"]][["p-value"]]))
  
  #embed.circaov from Rfast2 (originally from Directional)
  embed.circaovRfast2 <- apply(DistGroups,2,FUN=function(x) Rfast2::embed.circaov(x,as.factor(GroupsAB[,1])))
  embed.circaovRfast2p<-embed.circaovRfast2[2,]
  
 
  #hRfast2::het.circaov (originally from Directional)
  het.circaovRfast2 <- apply(DistGroups,2,FUN=function(x) Rfast2::het.circaov(x,as.factor(GroupsAB[,1])))
  het.circaovRfast2p<-het.circaovRfast2[2,]
  
  #hRfast2::het.circaov (originally from Directional)
  lr.circaovRfast2 <- apply(DistGroups,2,FUN=function(x) Rfast2::lr.circaov(x,as.factor(GroupsAB[,1])))
  lr.circaovRfast2p<-lr.circaovRfast2[2,]
  
  #Watson wheeler test! Test assumes von Mises
  watson.wheelerRES <- apply(DistGroups,2,FUN=function(x) watson.wheeler.test(x,GroupsAB[,1]))
  watson.wheelerP <- sapply(c(1:rans),function(x) watson.wheelerRES[[x]][["p.value"]])
  
  #Watson william test! Test assumes von Mises and equal kappa
  watson.williamsRES <- apply(DistGroups,2,FUN=function(x) watson.williams.test(x,GroupsAB[,1]))
  watson.williamsP<- sapply(c(1:rans),function(x) watson.williamsRES[[x]][["p.value"]])
  
  #Wallraff test!
  wallraff <- apply(DistGroups,2,FUN=function(x) wallraff.test(x,GroupsAB[,1],ref=NULL))
  wallraffP<- sapply(c(1:rans),function(x) wallraff[[x]][["p.value"]])
  
  ##MANOVA approach 
   funManCirc<-function(x) summary(manova(cbind(cos(x),sin(x)) ~ GroupsAB[,1]))
   MANOVA_approach <- apply(DistGroups,2, FUN=funManCirc)
   MANOVA_P <- sapply(c(1:rans),function(x) MANOVA_approach[[x]][["stats"]][1,6])
   
   #create a temp data.frame to store results
   temp <- data.frame(matrix(nrow = 1, ncol = 19))
   names(temp) <- c('tang.conc','levenes','WatsonTwoSamp','Watson.Wheeler',
                            "Watson.Williams",'Wallraff', 'Raodispersion', 
                            "Raopolar","ConcTestDirectional", "WatsonLargeSample",
                           "Fishers","FishersLarge", "MardiaWatsonWheeler", 
                            "Kuiper","Manova","P.Test.Assym","embed.circaovRfast2",
                    "het.circaovRfast2","lr.circaovRfast2")
   
  #####Calculate type I error/power in temp file 
  temp$levenes<- (sum(1*(LeveneP<0.05)))/rans
  temp$tang.conc<- (sum(1*(tangconcDirP<0.05)))/rans
  temp$WatsonTwoSamp<- (sum(1*(watson1>0.178)))/rans
  temp$Watson.Wheeler<- (sum(1*(watson.wheelerP<0.05)))/rans
  temp$Watson.Williams<- (sum(1*(watson.williamsP<0.05)))/rans
  temp$Wallraff<- (sum(1*(wallraffP<0.05)))/rans
  temp$Raodispersion <- sum(1*(raodispersionP == "Reject null hypothesis of equal dispersions "))/rans
  temp$Raopolar <- sum(1*(raopolarP != "Do not reject null hypothesis of equal polar vectors "))/rans
  temp$ConcTestDirectional<-(sum(1*(conctestDirP<0.05)))/rans
  temp$embed.circaovRfast2<-(sum(1*(embed.circaovRfast2p<0.05)))/rans
  temp$het.circaovRfast2<-(sum(1*(het.circaovRfast2p<0.05)))/rans
  temp$lr.circaovRfast2<-(sum(1*(lr.circaovRfast2p<0.05)))/rans
  temp$WatsonLargeSample<-(sum(1*(WatsonLargeP<0.05)))/rans
  temp$Fishers<-(sum(1*(FishersP<0.05)))/rans
  temp$FishersLarge<-(sum(1*(FishersLargeP<0.05)))/rans
  temp$MardiaWatsonWheeler<-(sum(1*(MardiaWatsonWheelerP<0.05)))/rans
  temp$Kuiper<- (sum(1*(kuiper1<0.05)))/rans
  temp$Manova<- (sum(1*(MANOVA_P<0.05)))/rans
  temp$P.Test.Assym<- (sum(1*(P.Test.Assym_P<0.05)))/rans

    temp # this is stored in  Power.sample
} #e

#the x-axis is calculated from the concentration of the second distribution, if the maximum is larger the minimum number
#otherwise it takes the mean direction vector
Power.sample$x=if(max(k2seq)>min(k2seq)) k2seq else deg(m2seq) 

Power.sample

#prepare the text of the csv. file 
text = paste(Name[i],"_n1_",nv1,"_n2_",nv2,"_its_",rans,"_Dist1_",typeDist1,"_Dist2_",typeDist2,
             "_m1=",deg(min(m1seq)),"-",deg(max(m1seq)),"_m2_",deg(min(m2seq)),"-",deg(max(m2seq)),
             "_k1=",min(k1seq),"-",max(k1seq),"_k2_",min(k2seq),"-",max(k2seq),".csv",sep="")

#write csv file in the working directory
write.csv(Power.sample, file = text,row.names=FALSE )

}#i
```
