## Supplementary figures for "Comparing two circular distributions: advice for effective implementation of statistical procedures in biology"

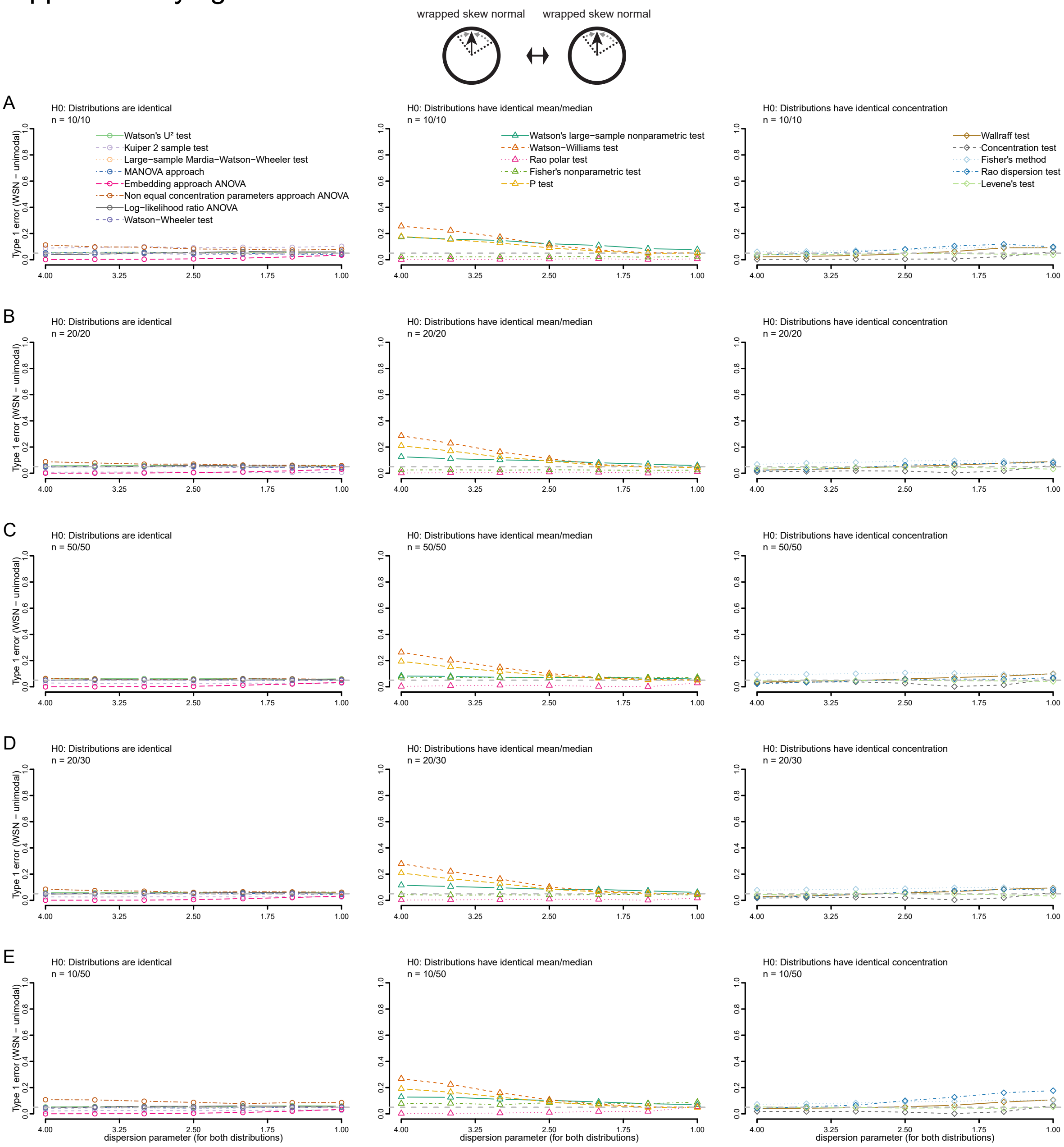


**Figure S1**: Type 1 error of all tests using wrapped skew normal distributions for different sample sizes: 10 and 10 (A), 20 and 20 (B), 50 and 50 (C), 20 and 30 (D) and 10 and 50 (E). The dispersion parameter decreases for both distributions from 4 to 1. Tests are grouped according to their null hypotheses.

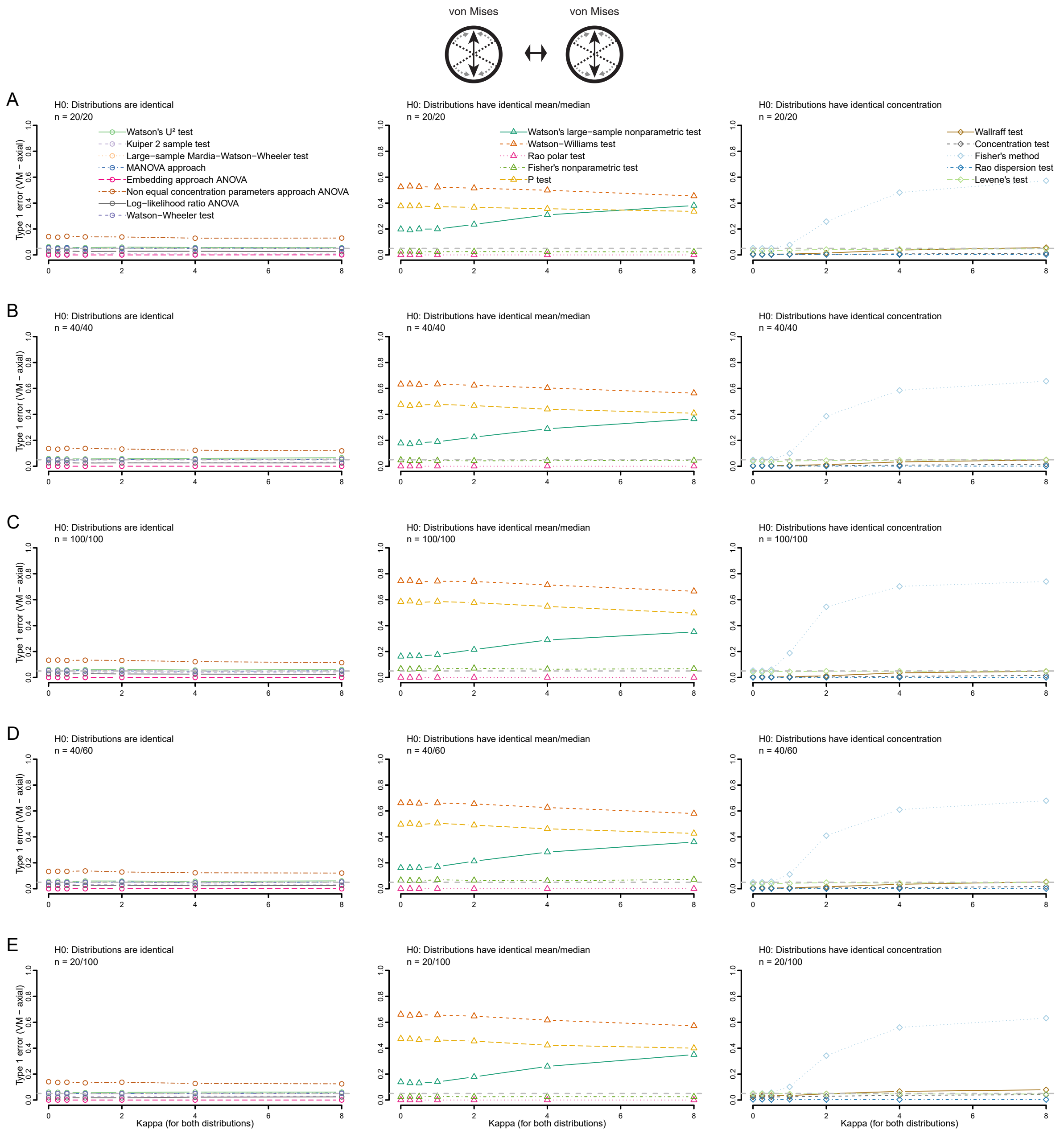


**Figure S2**: Type 1 error of all tests using axial von Mises distributions for different sample sizes: 20 and 20 (A), 40 and 40 (B), 100 and 100 (C), 40 and 60 (D) and 20 and 100 (E). The concentration parameter (κ, kappa) increases for both distributions from 0 to 8. Tests are grouped according to their null hypotheses.

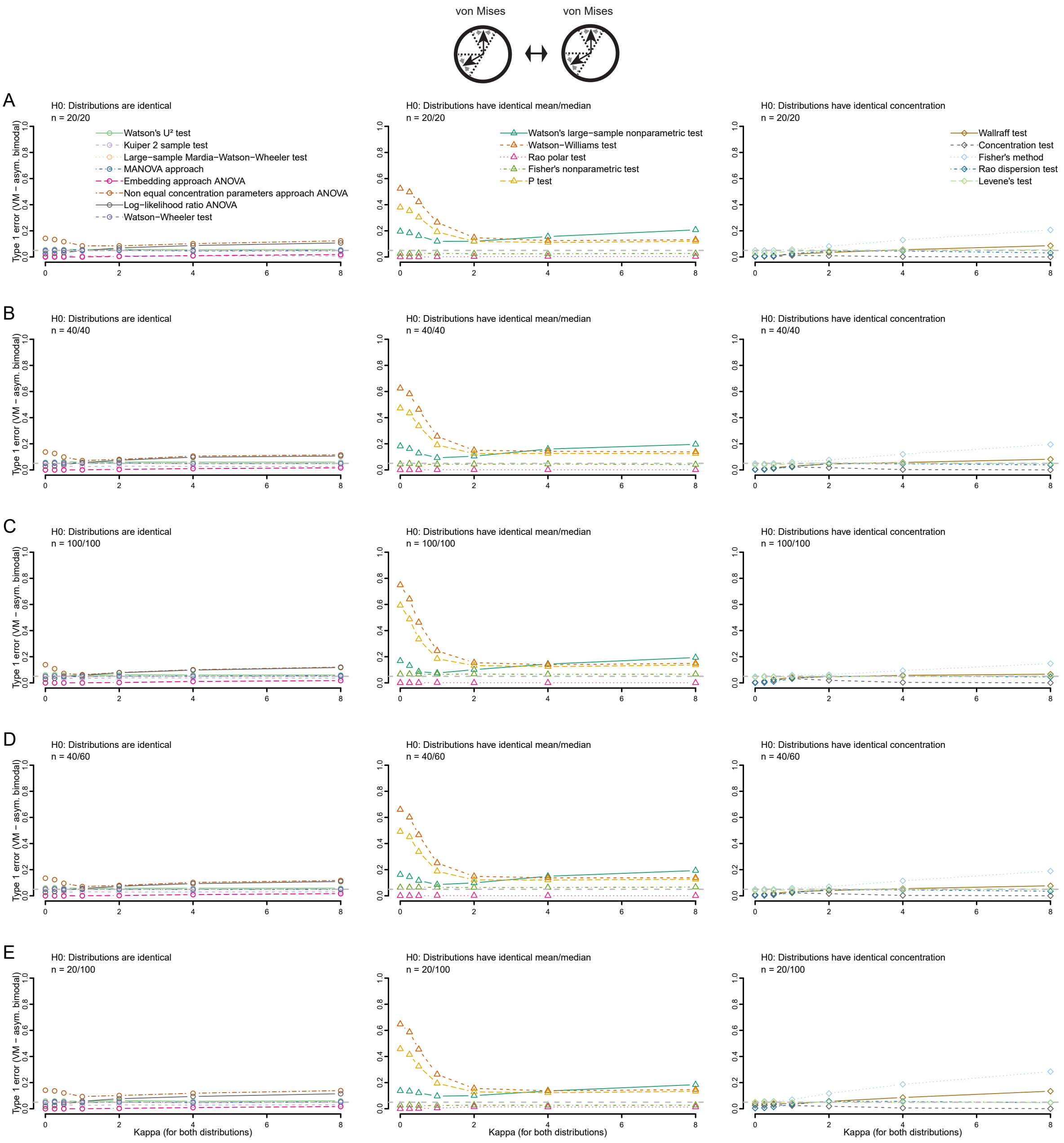


**Figure S3**: Type 1 error of all tests using asymmetrical bimodal von Mises distributions for different sample sizes: 20 and 20 (A), 40 and 40 (B), 100 and 100 (C), 40 and 60 (D) and 20 and 100 (E). The concentration parameter (κ, kappa) increases for both distributions from 0 to 8. Tests are grouped according to their null hypotheses.

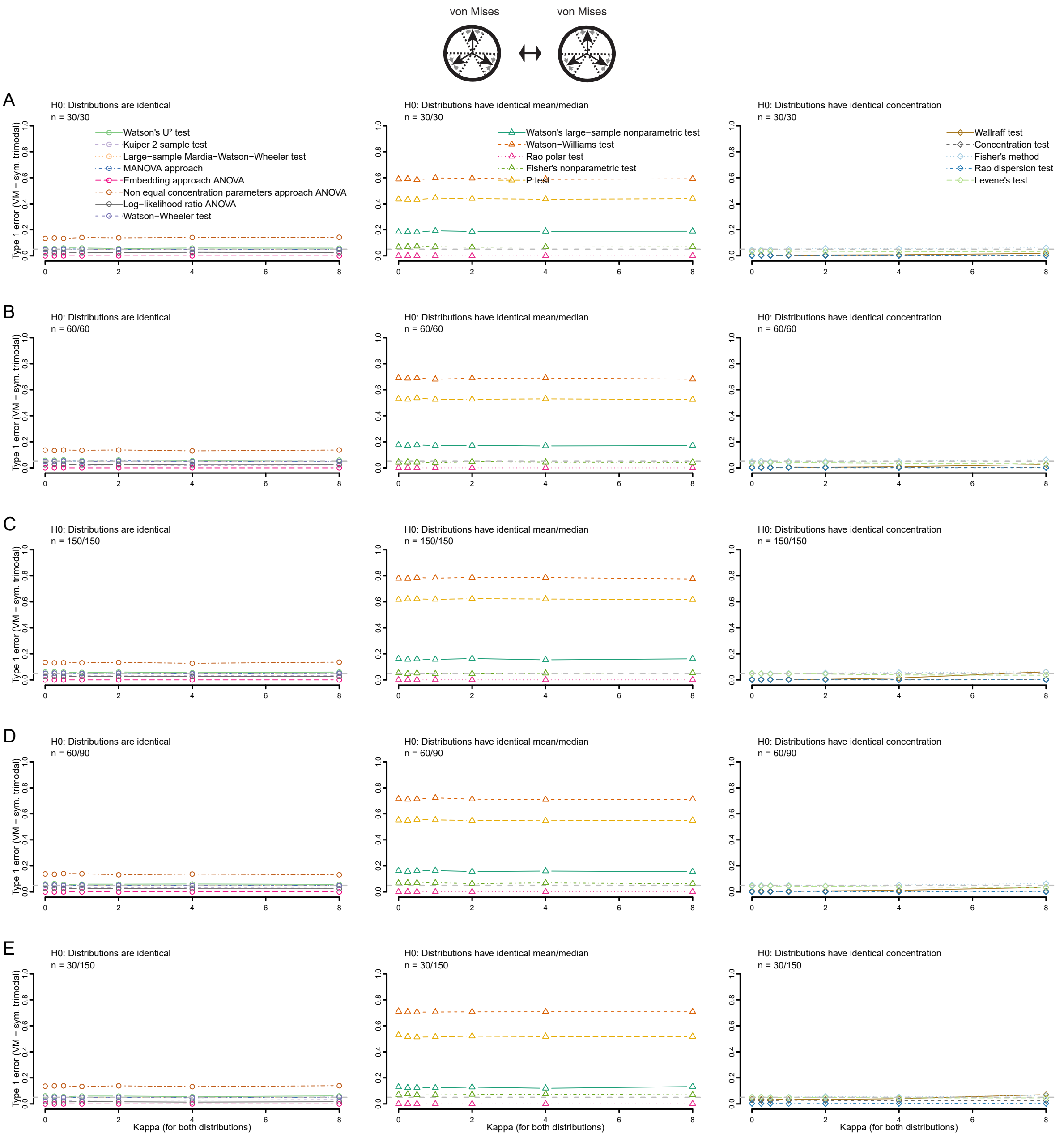


**Figure S4**: Type 1 error of all tests using symmetrical trimodal von Mises distributions for different sample sizes: 30 and 30 (A), 60 and 60 (B), 150 and 150 (C), 60 and 90 (D) and 30 and 150 (E). The concentration parameter (κ, kappa) increases for both distributions from 0 to 8. Tests are grouped according to their null hypotheses.

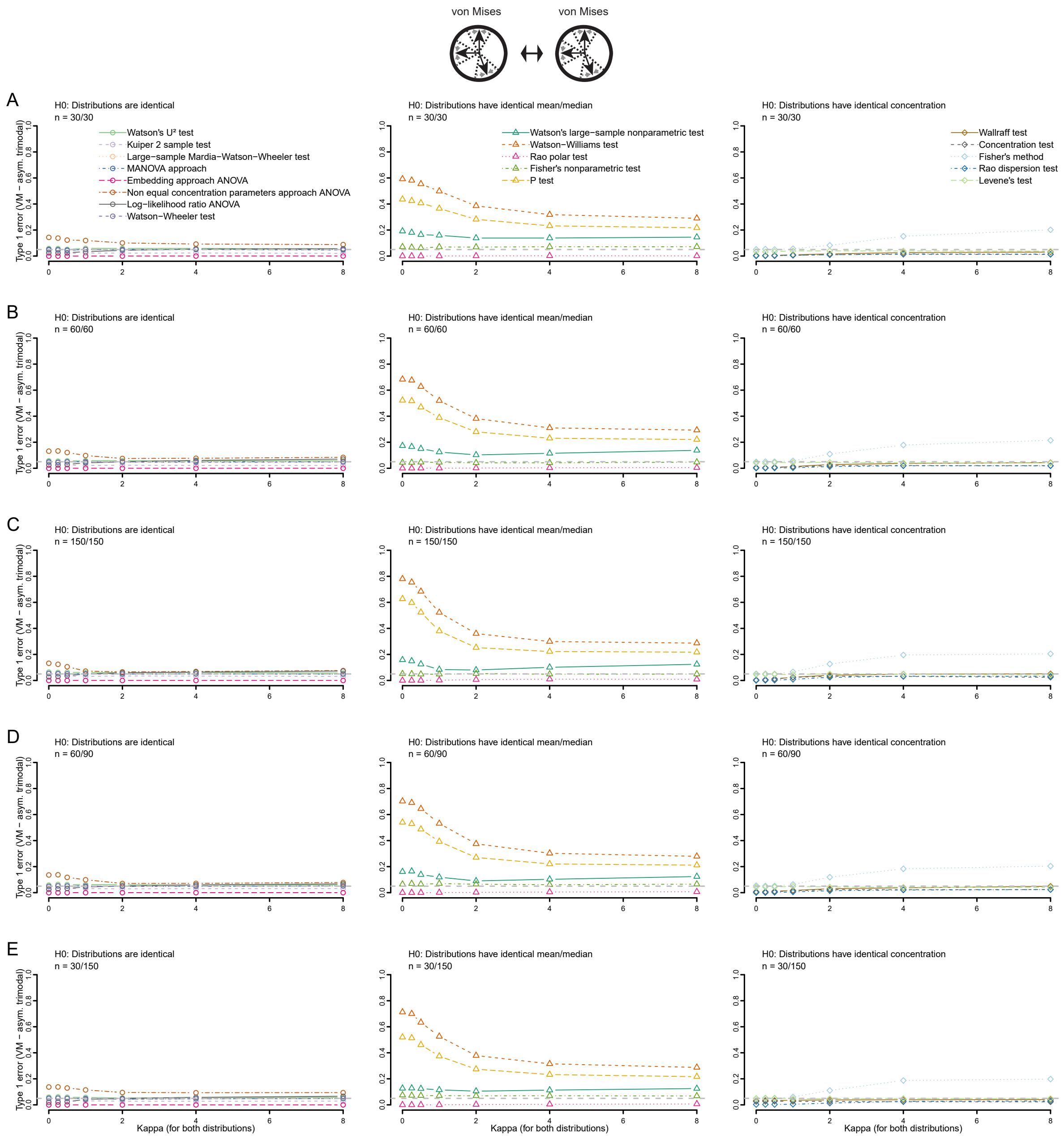


**Figure S5**: Type 1 error of all tests using asymmetrical trimodal von Mises distributions for different sample sizes: 30 and 30 (A), 60 and 60 (B), 150 and 150 (C), 60 and 90 (D) and 30 and 150 (E). The concentration parameter (κ, kappa) increases for both distributions from 0 to 8. Tests are grouped according to their null hypotheses.

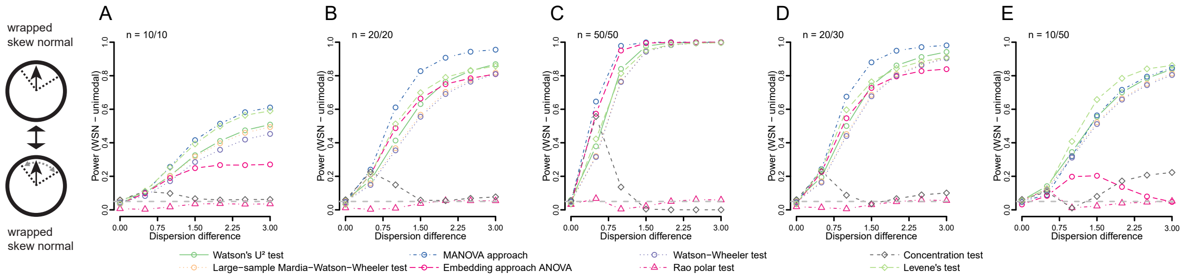


**Figure S6:** Power of all included tests when comparing wrapped skew normal distributions using different sample sizes: 10 and 10 (A), 20 and 20 (B), 50 and 50 (C), 20 and 30 (D) and 10 and 50 (E). The dispersion difference increases from 0 to 3.

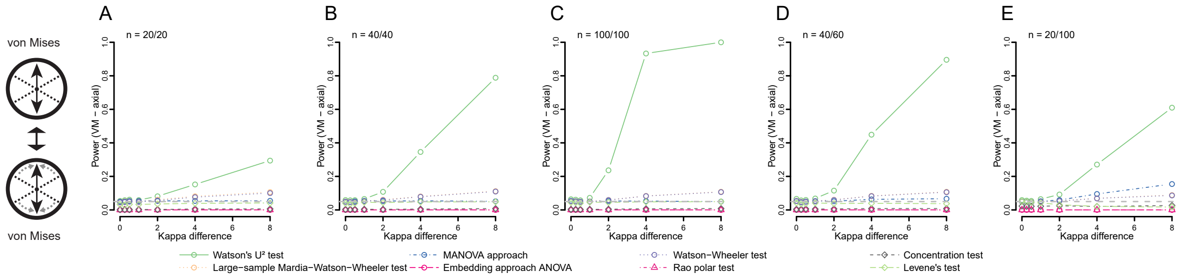


**Figure S7:** Power of all included tests when comparing axial von Mises distributions for different sample sizes: 20 and 20 (A), 40 and 40 (B), 100 and 100 (C), 40 and 60 (D) and 20 and 100 (E). The kappa difference increases from 0 to 8 (one distribution fixed at 0).

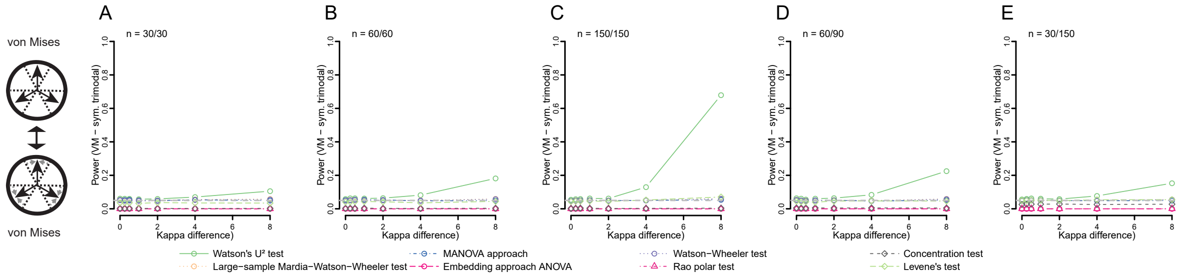


**Figure S8:** Power of all included tests when comparing symmetrical trimodal von Mises distributions for different sample sizes: 30 and 30 (A), 60 and 60 (B), 150 and 150 (C), 60 and 90 (D) and 30 and 150 (E). The kappa difference increases from 0 to 8 (one distribution fixed at 0).

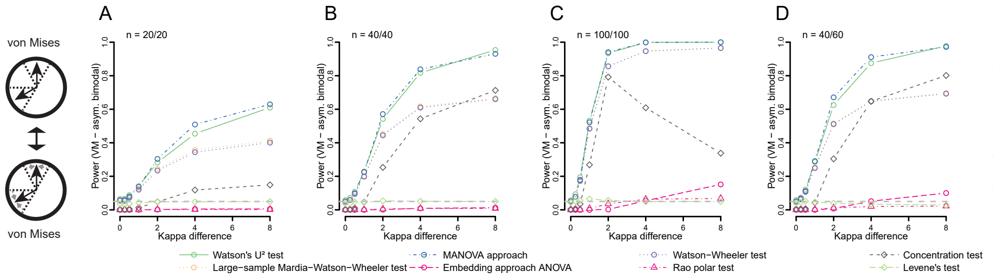


**Figure S9:** Power of all included tests when comparing asymmetrical bimodal von Mises distributions for different sample sizes: 20 and 20 (A), 40 and 40 (B), 100 and 100 (C), 40 and 60 (D) and 20 and 100 (E). The kappa difference increases from 0 to 8 (one distribution fixed at 0).

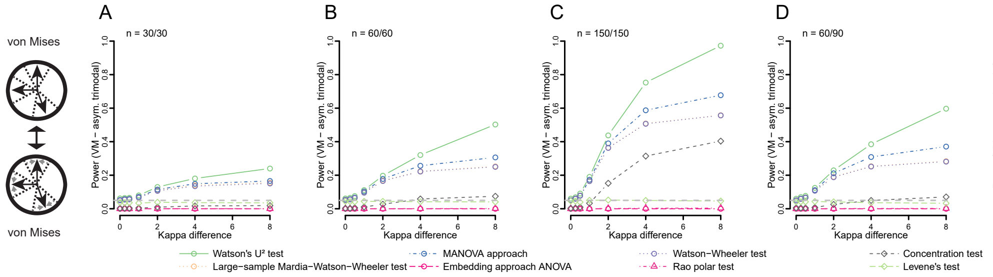


**Figure S10:** Power of all included tests when comparing asymmetrical trimodal von Mises distributions for different sample sizes: 30 and 30 (A), 60 and 60 (B), 150 and 150 (C), 60 and 90 (D) and 30 and 150 (E). The kappa difference increases from 0 to 8 (one distribution fixed at 0).

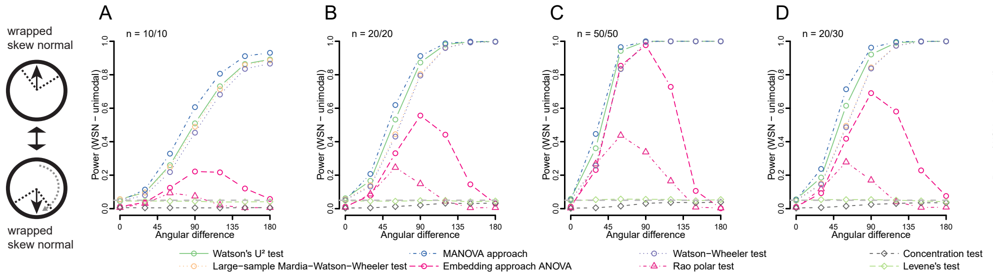


**Figure S11:** Power of all included tests when comparing wrapped skew normal distributions (dispersion parameter for both = 2) using different sample sizes: 10 and 10 (A), 20 and 20 (B), 50 and 50 (C), 20 and 30 (D) and 10 and 50 (E). The angular difference increases from 0 to 180 (one distribution fixed at 0).

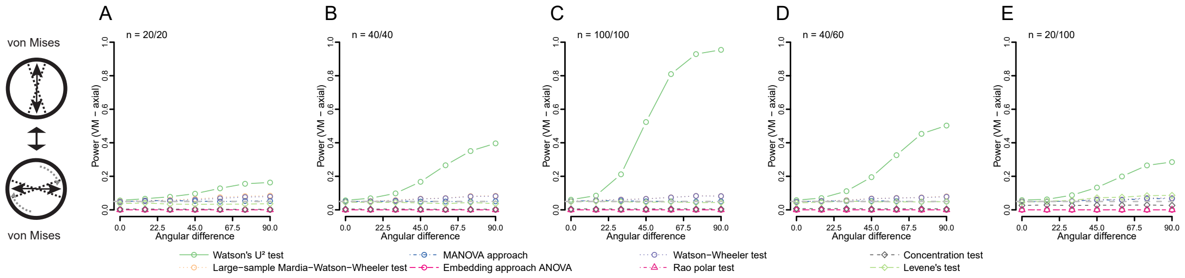


**Figure S12:** Power of all included tests when comparing axial von Mises distributions (kappa for both = 2) for different sample sizes: 20 and 20 (A), 40 and 40 (B), 100 and 100 (C), 40 and 60 (D) and 20 and 100 (E).The angular difference increases from 0 to 90 (one distribution fixed at 0).

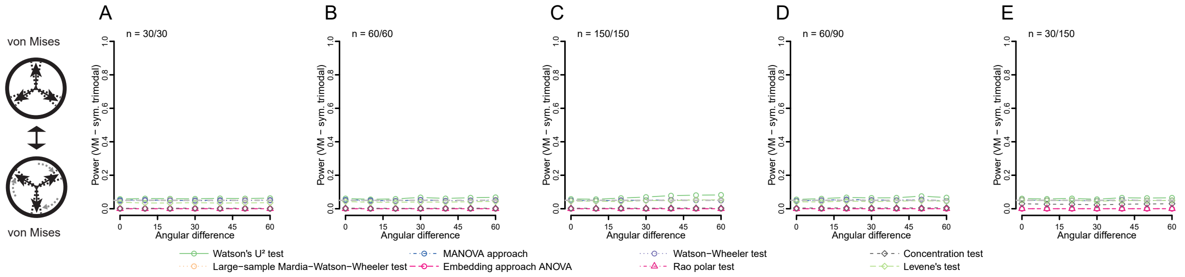


**Figure S13:** Power of all included tests when comparing symmetrical trimodal von Mises distributions (kappa for both = 2) for different sample sizes: 30 and 30 (A), 60 and 60 (B), 150 and 150 (C), 60 and 90 (D) and 30 and 150 (E). The angular difference increases from 0 to 60 (one distribution fixed at 0).

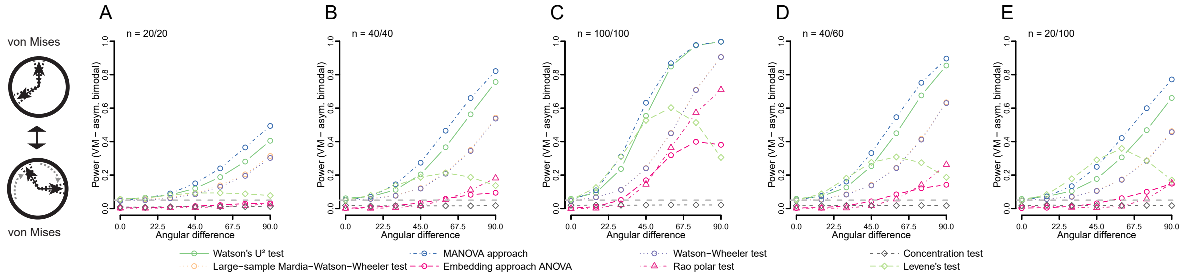


**Figure S14:** Power of all included tests when comparing asymmetrical bimodal von Mises distributions (kappa for both = 2) for different sample sizes: 20 and 20 (A), 40 and 40 (B), 100 and 100 (C), 40 and 60 (D) and 20 and 100 (E).The angular difference increases from 0 to 90 (one distribution fixed at 0).

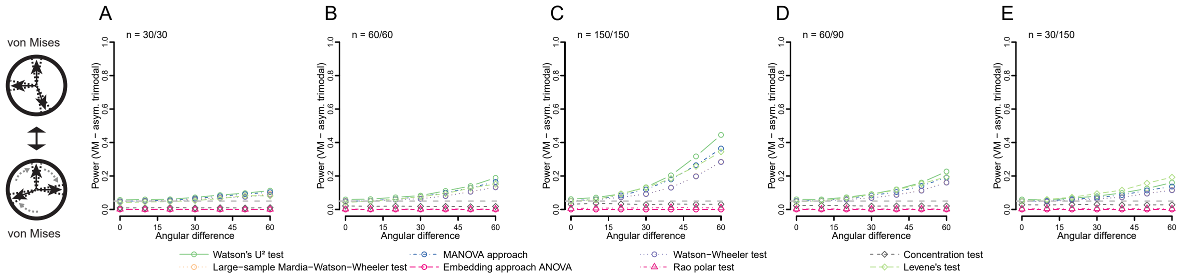


**Figure S15:** Power of all included tests when comparing asymmetrical trimodal von Mises distributions (kappa for both = 2) for different sample sizes: 30 and 30 (A), 60 and 60 (B), 150 and 150 (C), 60 and 90 (D) and 30 and 150 (E). The angular difference increases from 0 to 60 (one distribution fixed at 0).

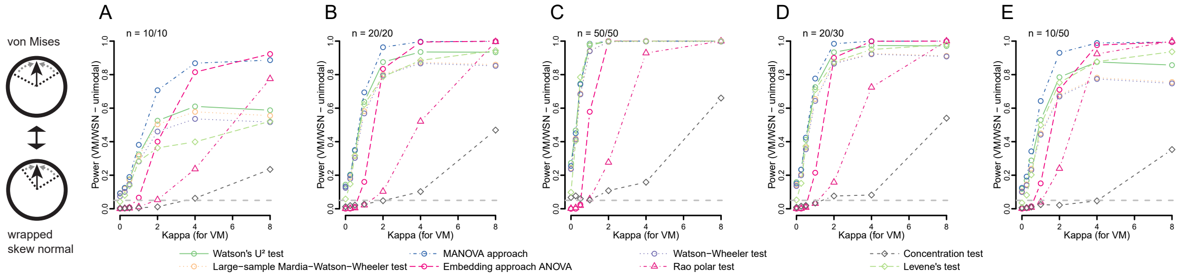


**Figure S16:** Power of all included tests when comparing von Mises with wrapped skew normal distributions using different sample sizes: 10 and 10 (A), 20 and 20 (B), 50 and 50 (C), 20 and 30 (D) and 10 and 50 (E). The concentration (κ) of the von Mises increases from 0 to 8, while the dispersion parameter of the wrapped skew normal decreases from 4 to 1.
